# Sexual selection and flight as predictors of sexual size and shape dimorphism in stick and leaf insects

**DOI:** 10.1101/2025.11.25.690473

**Authors:** Romain P. Boisseau, Sven Bradler, Douglas J. Emlen

## Abstract

Sexual size dimorphism (SSD) and sexual shape dimorphism (SShD) are widespread in animals, yet their macroevolutionary drivers remain poorly understood. We analyzed 196 stick and leaf insect species (Phasmatodea), a clade with exceptional diversity in size, shape, flight capacity, and mating systems, using phylogenetic comparative methods to evaluate the roles of fecundity selection, sexual selection, and ecological factors. SSD was universally female-biased and varied over an order of magnitude. Allometric scaling supported the inverse of Rensch’s rule, and clade-level patterns matched predictions from quantitative genetic models in which stronger directional selection on females than on males explains female-biased SSD. However, interspecific variation in SSD was not explained by female fecundity. Instead, mating system and flight dimorphism best predicted variation in SSD and SShD: males engaging in short-term mate guarding (less than a few days) were larger and stockier than searching males or those exhibiting prolonged mate guarding, reducing SSD, whereas species with flight-capable males and flightless females showed increased dimorphism, largely due to smaller males. Habitat and climate had limited effects. Sex differences in growth rate and development duration contributed equally to SSD, with females growing faster and for longer, consistent with widespread protandry. These results indicate that while fecundity selection historically drove female-biased SSD, contemporary variation is primarily shaped by male-specific selection through mating system and flight-related ecological pressures.

## Introduction

Phenotypic differences between the sexes are extremely widespread in animals and understanding their variation is a major question in evolutionary biology (Darwin 1871; Fairbairn 1997; Fairbairn et al. 2007). Body size differences in particular can be extreme (sexual size dimorphism; SSD), with mature females weighing hundreds of times more than their respective mates in some fishes, spiders, and marine invertebrates (Fairbairn 2013). Interspecific variation in the extent of SSD is hypothesized to reflect differences in the relative intensity of three types of sex-dependent selective pressures (Blanckenhorn 2005; Cox and Calsbeek 2009; Littleford-Colquhoun et al. 2019). First, fecundity selection is expected to favor larger female sizes when their fitness increases with the number and size of eggs produced (i.e., reproductive output), and when larger body sizes enable greater reproductive outputs (Shine 1988; Honěk 1993; Cox et al. 2003; Pincheira-Donoso and Hunt 2017). Second, sexual selection is expected to act primarily on male body size (Cox and Calsbeek 2009; Horne et al. 2020), ubiquitously favoring relatively larger male sizes in species with resource-defense or lek mating systems (e.g., through male-male combat or female choice) (Andersson 1994; Hardy and Briffa 2013; Emlen 2014) and sometimes favoring relatively smaller male sizes in species with scramble competition mating systems (e.g., through mobility selection) (Kelly et al. 2008; Herberstein et al. 2017). Finally, some aspects of natural selection may drive the evolution of sexual dimorphism through intersexual niche partitioning (Shine 1989). In such cases, sex differences may arise initially via sexual or fecundity selection and subsequently be exacerbated by sex-dependent natural selection and adaptation to separate niches (secondary ecological dimorphism hypothesis) (Fairbairn 1997; Stephens and Wiens 2009; Svenson et al. 2016). Understanding adaptive interspecific variation in SSD therefore requires quantifying the relative contributions of these three types of selective forces (i.e., sexual, fecundity and ecological).

SSD is usually accompanied by sexual dimorphism in shape. Males and females may differ in body proportions (Butler et al. 2007), the shape of body structures (Berns and Adams 2013), ornaments and weaponry (Emlen 2008) and coloration (Dale et al. 2015). Interspecific variation in sexual shape dimorphism (SShD) is thought to result from the same three sex-dependent selective pressures (Adams et al. 2020), with fecundity selection primarily affecting female shape, sexual selection primarily affecting male shape, and some aspects of ecological selection acting differentially on both male and female shape.

We explored the selective drivers of interspecific variation in SSD and SShD across the stick and leaf insects (order Phasmatodea). Specifically, we tested whether (I) variation in fecundity selection was more strongly associated with variation in female than male size or shape; (II) variation in sexual selection was more strongly associated with variation in male than female size or shape; and (III) variation in ecological selection was differentially associated with variation in female or male size or shape.

Phasmids exhibit a spectacular diversity of body forms and sizes ranging from very elongated to much more robust bodies and from giant to more modest sizes (Bedford 1978; Bradler and Buckley 2018; Brock and Büscher 2022). For instance, the branch-like female *Phryganistria chinensis* (Zhao, informal name), considered the longest insect in the world, measures up to 37cm long (excluding legs and antennae)(Shi et al. 2019) while the sturdy female *Miniphasma prima* (Zompro, 1999) only grows to 1.7cm (Zompro 1999). This diversity of body forms likely stems from the advergence of phasmid appearance towards various objects of their habitat (e.g., sticks, leaves, bark pieces, moss) which enables an astonishing camouflage through crypsis and masquerade (Boisseau et al. 2025). Even more notably, sexual dimorphism can be very strong in this group, to the point of confusing taxonomists into assigning males and females of the same species to separate genera (Cumming et al. 2020). As is most common in arthropods, males are usually (much) smaller and slenderer than females (Sivinski 1979). In addition, specific traits such as relative leg size and shape, coloration, or the presence of wings or ocelli (i.e., simple eyes) are often dimorphic (Boisseau et al. 2020; Bank and Bradler 2022). In many species, the much larger and sedentary females are flightless, either with shortened wings or no wings at all, while the mobile males can be fully winged and flight capable (Zeng et al. 2020, 2023). Phasmids also exhibit a remarkable array of reproductive modes and mating systems, from small mobile males searching for scattered sedentary females (Boisseau et al. 2022), to tiny monogamous males riding on the back of a female for life (Conle et al. 2009), large weapon-bearing males aggressively fighting for territories and females (Boisseau et al. 2020; Boisseau and Emlen 2025), and sometimes all the way to obligate thelytokous parthenogenesis (Schwander and Crespi 2009; Schwander et al. 2025). Therefore, stick and leaf insects constitute an ideal group in which to investigate the selective drivers of both SSD and SShD.

In the present study, we used morphological data on 196 phasmid species (6% of described species diversity and 27% described generic diversity) to uncover patterns of body size, shape, and wing sexual dimorphism in the group. We first mapped and quantified the variation of SSD and SShD across species. Then we tested for effects of female lifetime reproductive output (a proxy for fecundity), mating system (a proxy for the direction and strength of sexual selection), and sex-specific flight capacity, climate, and habitat (proxies for dimorphic ecological selection) on the variation of each sex’s size and shape, and on SSD and SShD. Finally, we also investigated proximate mechanisms contributing to SSD in this group by evaluating the relative importance of sex differences in growth and development time.

## Materials and methods

The present study was carried out in tandem with a companion study focusing on ecomorphological convergence in female morphology across Phasmatodea (Boisseau et al. 2025). As such, the two studies used the same phylogeny, methods and data on female morphology and habitat classification.

### Taxon sampling and phylogeny

We used a phylogeny including a total of 314 phasmid taxa (9% of phasmid species diversity and 33% of generic diversity) and one embiopteran species as outgroup. The phylogenetic reconstruction was carried out using genetic data from 3 nuclear (*i.e.,* 18S rRNA, 28S rRNA and histone subunit 3) and 4 mitochondrial genes (*i.e.,* 12S rRNA, 16S rRNA, cytochrome-c oxidase subunit I and II), the basal topology of transcriptome-based trees (Simon et al. 2019; Tihelka et al. 2020) and Bayesian inferences (see Boisseau et al. 2025 for details).

### Morphological data

We measured 1,365 adult female (212 species) and 1,000 adult male specimens (200 species, 196 species with specimens of both sexes). Measurements from female specimens have already been used in the companion study focusing on female morphology (Boisseau et al. 2025). We used ImageJ (v1.51) (Schneider et al. 2012) and high quality photographs of live or pinned specimens in dorsal and/or lateral view from our own collection at the University of Göttingen (Germany), field guides (Brock and Hasenpusch 2009; Seow-Choen 2016, 2017, 2018), the published literature, and various online databases (Brock et al. 2021) (Dataset S1). Between 1 and 23 different individuals per sex and per species (mean= 6.4 females and 5 males per species) were measured. Because photographs were sometimes unscaled, we measured each of the 23 continuous traits described in Figure S1 relative to body length. We measured body length directly (excluding ovipositor and subgenital plate) on properly scaled photographs and collected body length data from the literature (Dataset S1). The median of each relative measurement was then multiplied by the sex- and species-specific median of body length to obtain absolute measurements for each sex and species. Further methodological details are available in (Boisseau et al. 2025).

Body volume was used as a proxy for body size and corresponded to the volume of an elliptic cylinder calculated from body length, average body width and average body height (Figure S1). SSD was quantified using a commonly used sexual dimorphism index (SDI) (Lovich and Gibbons 1992):

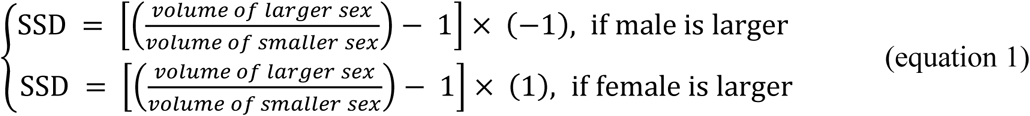

Females were always the larger sex in our dataset so SSD was always positive. To quantify body shape while controlling for size, we performed a Principal Component Analysis (PCA, *R function*: “R package”; *prcomp*: “stats”) including all absolute measurements (except body volume), but original values were substituted with residuals calculated from a phylogenetically-corrected generalized least-squares (PGLS) regression against body volume (*pgls*: “caper” (Orme et al. 2018), lambda=’ML’). Wing length and area, which included zeros for wingless species, were divided by body length or body length squared respectively to obtain measures of relative wing length and area. These variables were all mean-centered and scaled to unit variance. PC1 accounted for 59% of the total variation and reflected body elongation, while PC2 accounted for 12% of the total variation and reflected wing size (Figure S2). We focused on these two aspects of shape to calculate SShD in body elongation (SShD_elongation_) and relative wing size (SShD_wings_) as the difference between male and female PC1 or PC2 respectively. A larger and positive SShD_elongation_ and SShD_wings_ respectively indicate that males are more elongated than females, or have larger wings. We also calculated overall SShD as the Euclidean distance between males and females considering their coordinates along PC1-PC4 (cumulatively accounting for 86% of the variation).

### Life history and ecological data

Stick insects are elusive, nocturnal animals whose natural history in the wild is poorly known. However, many species are bred in captivity as pets, especially in Europe. Thus, some information about their natural history and behavior is available from amateur enthusiasts and breeders. We surveyed the literature, field guides, breeding guides and online databases to collect information about fecundity, mating system, flight capacities and habitat (Dataset S1).

#### Fecundity

Species-level data on egg volume (proxy for egg size) and female lifetime fecundity (in number of eggs) were available from a companion study (Boisseau and Woods 2024). Female fecundity was quantified as their lifetime reproductive output, calculated as the product of egg volume and the lifetime number of eggs laid (n=90 species with sufficient data). Under the hypothesis that variation in fecundity selection (i.e., favoring larger females) drives variation in SSD and SShD, we predicted that species where females have a relatively higher reproductive output would display a relatively higher SSD and body SShD.

#### Mating system

We classified mating systems of 101 species into three broad categories: “searching”, “guarding”, and “monogamy” based on the typical duration of the association between males and females, including the initial copulation and any post-mating guarding period. Males of species in the “searching” category typically spend less than a day paired up with a single female. In these species, male fitness is probably mostly determined by their capacity to locate and move efficiently between scattered females, not by direct male-male competition. In this context, we expect smaller and more mobile males to be favored as they likely benefit from a more efficient locomotion (walking or flying) (Kelly et al. 2008; Boisseau et al. 2020, 2022).

Males of species in the “guarding” category typically stay with the same female for more than a day after finding it. Intromission may occur only initially or be repeated intermittently. Either way, the male often spends most of his time mounted on the female but not actively transferring sperm. Males eventually leave the female after a few days, typically less than four days, to search for another mate. This mate guarding strategy suggests a higher risk of sperm competition and a higher level of sexual selection, and therefore male fitness may also depend on the ability of a male to defend and guard access to a female from rival males. Direct combats between a guarding and an intruder male have been reported in several species (Sivinski 1979; Boisseau et al. 2020; Boisseau and Emlen 2025; Wilner et al. 2025). In this case, we expect both male fighting and mobility abilities to be important. Given that, unlike mobility selection, direct male combat almost ubiquitously favors larger sizes, we expect male size in these species to reflect this balance and to be relatively larger than species with non-guarding males.

Finally, some phasmid males spend more than four days, and in this case often their lifetime, riding on one female’s back, often staying in copula with their body essentially acting as a sperm plug (Conle et al. 2006). In these largely monogamous species, we expect male and female fitness interests to align and therefore smaller male sizes to be favored so as not to impair female locomotion (walking or flying), and to limit their need to forage (potentially away from the female).

#### Climatic data

We gathered information about the geographic range of each species based on sampling location of type specimens and observations on iNaturalist (available from https://www.inaturalist.org. Accessed July 2021). For each species, we then selected the median location with the most central latitude. From the GPS coordinates of the most central location for each species, we extracted data on annual mean temperature, mean diurnal range (i.e., mean of monthly (maximum - minimum temperature)), temperature seasonality (i.e., standard deviation ×100), annual temperature range (i.e., maximum temperature of warmest month - minimum temperature of coldest month), annual precipitation, precipitation seasonality (i.e., coefficient of variation) from worldclim (Fick and Hijmans 2017) (available from https://www.worldclim.org. Accessed July 2021). We also extracted the length of the growing period (i.e., number of days during a year when temperatures are above 5°C and precipitation exceeds half the potential evapotranspiration (van Velthuizen et al. 2007), available from https://data.apps.fao.org/map/catalog), net primary production of biomass (grams of dry matter per m^2^ per year; Climate Research Unit, Univ. of East Anglia, period 1976-2000, available from https://data.apps.fao.org/map/catalog) from the FAO Map Catalog; and total annual growing degree days (i.e., a measure of the annual amount of thermal energy available for plant and insect growth; Climate Research Unit, Univ. of East Anglia, available from https://sage.nelson.wisc.edu/data-and-models). Such data was available for all species (n=196). Given that these nine climatic variables are highly correlated, we ran a principal component analysis (Figure S3). We kept the first component (climate PC1, explaining 60.5% of the total variation) to quantify climatic variation between species. Climate PC1 mainly reflected net primary productivity (i.e., food availability for herbivorous insects) and was positively correlated with annual precipitation and mean temperature, and negatively correlated with the annual and diurnal temperature range (i.e., seasonality). Thus, PC1 is overall high in tropical regions, and low in more temperate and seasonal regions. Season length and food availability may limit the growth of some species. Because females are larger than males, we expect this hypothetical growth limitation to affect primarily females and therefore to reduce SSD. Therefore we predicted a higher SSD in tropical regions (high PC1, long growing season) than in temperate regions.

#### Flight dimorphism

We recently showed that large body sizes impair the flying performance of male phasmids (Boisseau et al. 2022). Larger males, with a relatively high wing loading, have more difficulties climbing and maneuvering in the air and are at higher risk of slipping and crashing when landing. This suggests that the mode of locomotion of males and females –i.e., flying and walking, or walking only—is likely to affect their size and shape evolution. We gathered observational data on the flight capacities of both sexes across species. Each sex was scored as either flight capable (including gliding) or flightless (including parachuting). Because flight can be reliably predicted from morphological characteristics and especially the size of the wings relative to the size of the animal (Zeng et al. 2023), we trained random forest models (i.e., a classification machine learning algorithm) to predict flight in species with missing observations (*randomForest*: “randomforest” (Liaw and Wiener 2002)). The algorithm was trained separately for each sex on the entire subset of the dataset for which flight data could be collected and was provided with data on sex-specific body volume, body length, relative body width, femur length, wing length, wing aspect ratio and wing area. Error rate was 1.18% for males and 2.16% for females. We used the trained algorithms to predict both male and female flight in species with missing flight data (n=8 species for female and n=19 species for male flight, over 196 species total). Finally, we qualitatively categorized flight dimorphism in each species as either both sexes being flightless, flight capable, or males being flight capable and females flightless. We expected species with only males able to fly to be more dimorphic as ecological selection related to flight would only affect males and favor relatively smaller sizes with larger wings.

*Habitat.* Finally, we classified each species’ habitat as the vegetation layer (i.e., ground/shrub, shrub/understory and understory/canopy) that they typically inhabit during the daytime (i.e., when they are exposed to visually hunting predators). The ground/shrub layer includes species mostly found resting below 1.5m (i.e., in the leaf litter, in the grass, in low shrubs, on logs, etc.). The shrub/understory layer corresponds to species resting at heights between 1.5m and 4m (i.e., in tall shrubs, on tree trunks, or in small trees). Finally, the understory/canopy layer corresponds to species typically resting in the canopy of tall trees (>4m). Habitat could be classified for 195 species. These strata differ in their structural and aerodynamic properties, predator communities and visual characteristics. For instance, unlike the ground/shrub layer, the canopy is characterized by substantial gaps, wind gusts, and a visual environment dominated by green leaves. Males and females often differ in their locomotor needs: males need to move to find females while females may only need to move to feed and sometimes to lay eggs. Thus, we expect that the vegetation layer where a species mainly resides may affect sexual dimorphism through ecological selection on locomotor performance. We specifically predicted that higher strata would be associated with higher levels of sexual dimorphism as locomotion may be more challenging in high gappy habitats.

### Ontogenetic data

We collected data on sex-specific postembryonic development duration (from hatching to final molt) from the literature and amateur breeding guides (Dataset S1). Despite such information often being reported qualitatively (± 2 weeks) without accounting for temperature, most phasmids are captive bred at room temperature (19-24°C) and exhibit extensive variation in development time across species (from 2 to 13.5 months) relative to measurement error, thus rendering the data informative. Sex-specific relative growth rate (RGR) was calculated as:

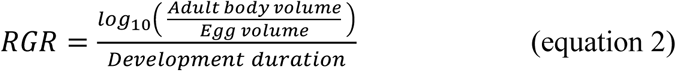

RGR therefore represents a growth rate averaged over the entire postembryonic development period (Tammaru and Esperk 2007). Egg volume was used as a proxy for the volume of first-instar individuals, assuming that egg sizes do not differ between males and females, which has, to our knowledge, never been observed in phasmids. Development duration and RGR could be found or calculated in both sexes in a total of 67 species.

Because females always had a higher (or equal) RGR and development time than their respective males, we calculated sex differences in RGR (SDRGR) and duration of development (SDDD) as:

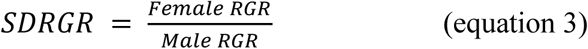

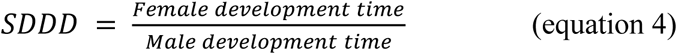

### Statistical analyses

Statistical analysis were carried out in R (v4.1.1) (R Core Team 2021). SSD and overall SShD were log_10_-transformed to improve data distribution for the subsequent analyses. We first mapped the evolution of SSD and overall SShD on the Phasmatodean phylogeny using a fast estimation of maximum likelihood ancestral states (*fastAnc*: “phytools” (Revell 2012)).

#### Allometry and SSD

Female-biased SSD in phasmids may have evolved primarily through negative directional selection acting on male size or positive directional selection on female size. Quantitative genetic theory predicts that the sex under historically stronger directional selection will exhibit greater interspecific variation. Consequently, as the extent of SSD and the allometry between log_10_ male versus log_10_ female body size reflects the history of selection on male and female size, we expect a correlation between the interspecific allometric slope (𝛽) of log_10_ male versus log_10_ female body volume and the extent of SSD across related clades (Zeng 1988; De Lisle and Rowe 2013). A positive correlation between 𝛽 and SSD among related clades would indicate that selection on average has acted more intensely on male body size, while a negative correlation would indicate more intense selection on female size as the primary explanation for why females are larger than males in phasmids. Therefore, for each phasmid clade with sufficient data (n_species_ ≥ 6), we calculated 𝛽 both as the phylogenetic reduced major axis (PRMA, *phyl.RMA*: “phytools”) and phylogenetic generalized least-squares (PGLS, *gls*: “nlme”, *corPagel*: “ape” (Paradis and Schliep 2019), Pagel’s 𝜆 estimated by maximum likelihood) regression slopes of log_10_ male versus log_10_ female body volume. A phylogenetically controlled mean SSD was also computed for each clade by using an intercept-only PGLS model in which log_10_ SSD was the response variable. Pharnaciini, the European clade, Stephanacridini and Agathemeridae (comprising less than 5 species included in the phylogeny) were grouped with their closest relatives, respectively Clitumninae + Pachymorphinae, the African/Malagasy clade, Lanceocercata and Pseudophasmatinae.

The correlation between 𝛽 and mean SSD across major clades was tested using a PGLS model taking into account the phylogenetic relationships between the clades. Similarly, Rensch’s rule suggests that SSD will be negatively correlated with body size (average between males and females) in species in which females are larger than males, and that males are always the sex with the greatest interspecific variation in size (i.e., 𝛽 > 1) (Rensch 1959). To test Rensch’s rule across Phasmatodea, we first quantified the allometry of log_10_ male versus log_10_ female body volume across all phasmid species, using both a PGLS and PRMA, as advised in a recent study (Meiri and Liang 2021). We also regressed log_10_ SSD on the mean body volume of males and females of each species using a PGLS.

#### Proximate causes of SSD

Females may be larger than males because they develop faster and/or for longer. We tested the association between SSD and sex difference in duration of post embryonic development and sex difference in relative growth rate using PGLS models. We tested the effect of log_10_ (sex difference in duration of development) and log_10_ (sex difference in relative growth rate) (explanatory variables) on log_10_ SSD (response) using two single predictor and one multiple predictor model. log_10_ (sex difference in duration of development) and log_10_ (sex difference in relative growth rate) were mean-centered on zero and scaled to unit variance to allow meaningful comparison of their effect sizes and relative importance in determining SSD. In the multiple predictor model, the significance of the effect of each explanatory variable was assessed using a type III ANOVA.

#### Fecundity, sexual selection, and ecological selection drivers of SSD and SShD

To evaluate the predictions from the non-mutually exclusive ultimate hypotheses I-III laid out in the introduction, we tested the effect of variation in ecological variables and reproductive life-history characteristics on the variation of female and male body volume, body elongation (PC1) and relative wing size (PC2), and consequently on the extent of SSD, SShD_elongation_ and SShD_wings_. We used either log_10_ male or female body volume, body elongation (PC1) or relative wing size (PC2), or log_10_ SSD, SShD_elongation_ and SShD_wings_ as dependent variables and log_10_ lifetime female reproductive output (i.e., fecundity), climate PC1, mating system, flight dimorphism and habitat as explanatory variables in single predictor PGLS models (including each explanatory variable alone to maximize sample size) and in multiple predictor PGLS models (including all five explanatory variables at once and no interaction). All continuous variables were mean-centered and scaled to unit variance to enable meaningful comparisons of effect sizes. In multiple predictor models, the significance of the effect of each explanatory variable was assessed using type II ANCOVAs. For categorical predictors showing significant effects, we subsequently performed post-hoc pairwise comparisons of estimated marginal means between groups using the Holm correction to account for multiple testing (*emmeans*:”emmeans”). We evaluated the proportion of variance explained by the PGLS models by calculating two R^2^ coefficients (*R2.resid*: “rr2” (Ives and Li 2018)): R^2^ , the total variance explained by the full model, and R^2^ , the variance explained by the fixed effects only, after accounting for phylogeny.

## Results

### Variation in SSD and SShD across Phasmatodea

SSD and SShD varied extensively across clades and species of stick and leaf insects (Figure 1-2, Table S1). SSD was exclusively female-biased but varied from females only being 1.25 times more voluminous than males in *Epicharmus marchali* (Lanceocercata), all the way to 22.5 times larger in *Cranidium gibbosum* (Diapheromerinae). SSD and overall SShD were loosely but significantly correlated (Figure S4; PGLS: 𝜆 = 0.59, F_1,184_ = 9.82, p = 0.002, R^2^ = 0.05). Thus, strongly size dimorphic species tended to also exhibit strongly dimorphic body shapes, especially with much smaller males also being much slenderer and carrying larger wings.

**Figure 1:**
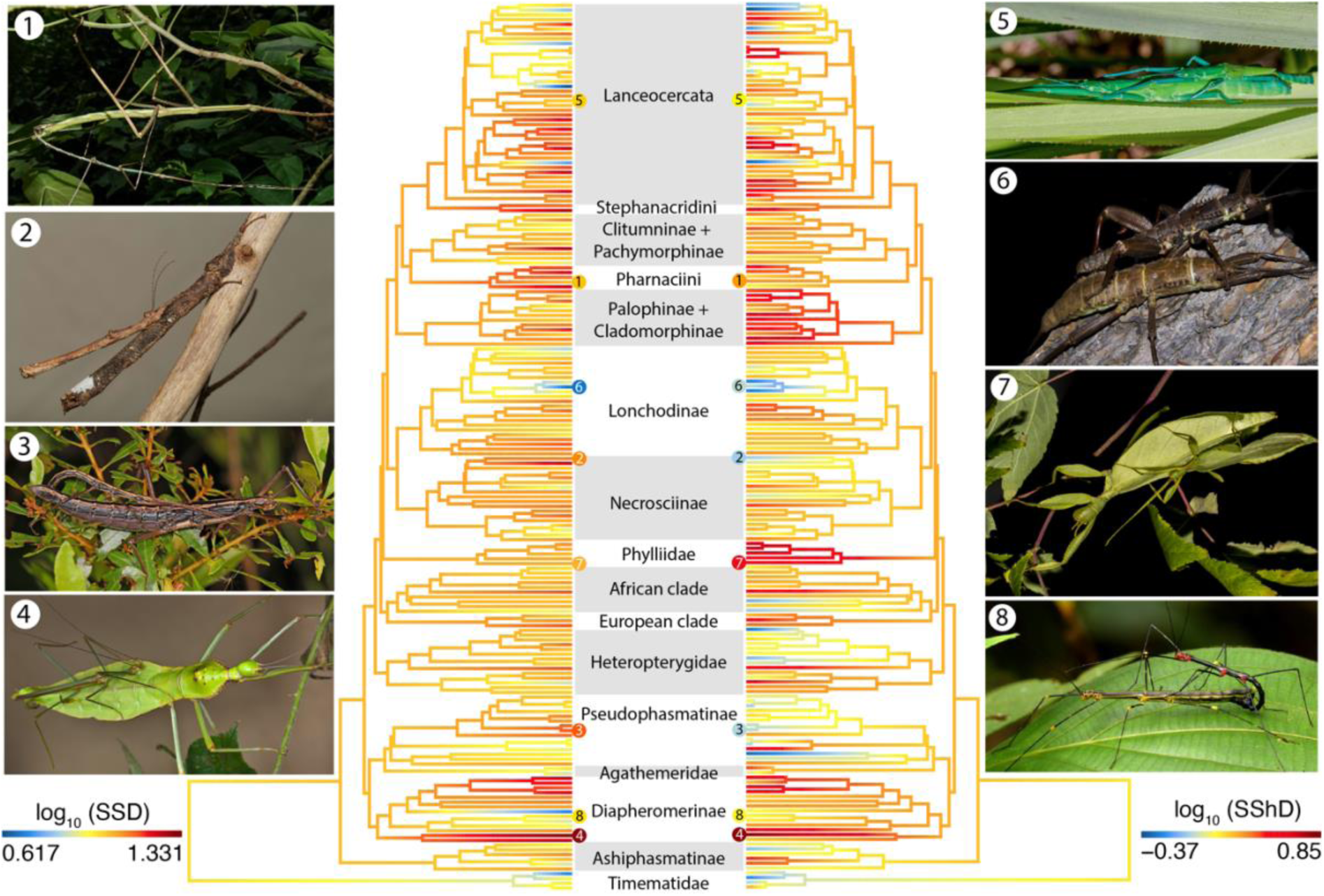
Diversity and phylogenetic pattern of sexual size dimorphism (SSD) and sexual shape dimorphism (SShD) in stick and leaf insects. The trees show the ancestral state reconstructions of SSD and overall SShD on the Phasmatodean tree. Numbered tips correspond to the pictures of mating pairs on the sides. **Picture 1**: *Phryganistria sp.* (Pharnaciini), photo by Rejoice Gassah (CC BY 4.0). **Picture 2**: *Neoclides buesheri* (Necrosciinae), photo by Bruno Kneubühler (used by permission). **Picture 3**: *Anisomorpha buprestoides* (Pseudophasmatinae), photo by Judy Gallagher (CC BY 2.0). **Picture 4**: *Cranidium gibbosum* (Diapheromerinae), photo by Bruno Kneubühler (used by permission). **Picture 5**: *Megacrania batesii* (Lanceocercata), photo by David White (CC BY-NC 4.0). **Picture 6**: *Eurycantha calcarata* (Lonchodinae), photo by Romain Boisseau. **Picture 7**: *Phyllium philippinicum* (Phyllidae), photo by Romain Boisseau. **Picture 8**: *Oreophoetes topoense* (Diapheromerinae), photo by Andreas Kay (CC BY-NC-SA 2.0).

**Figure 2:**
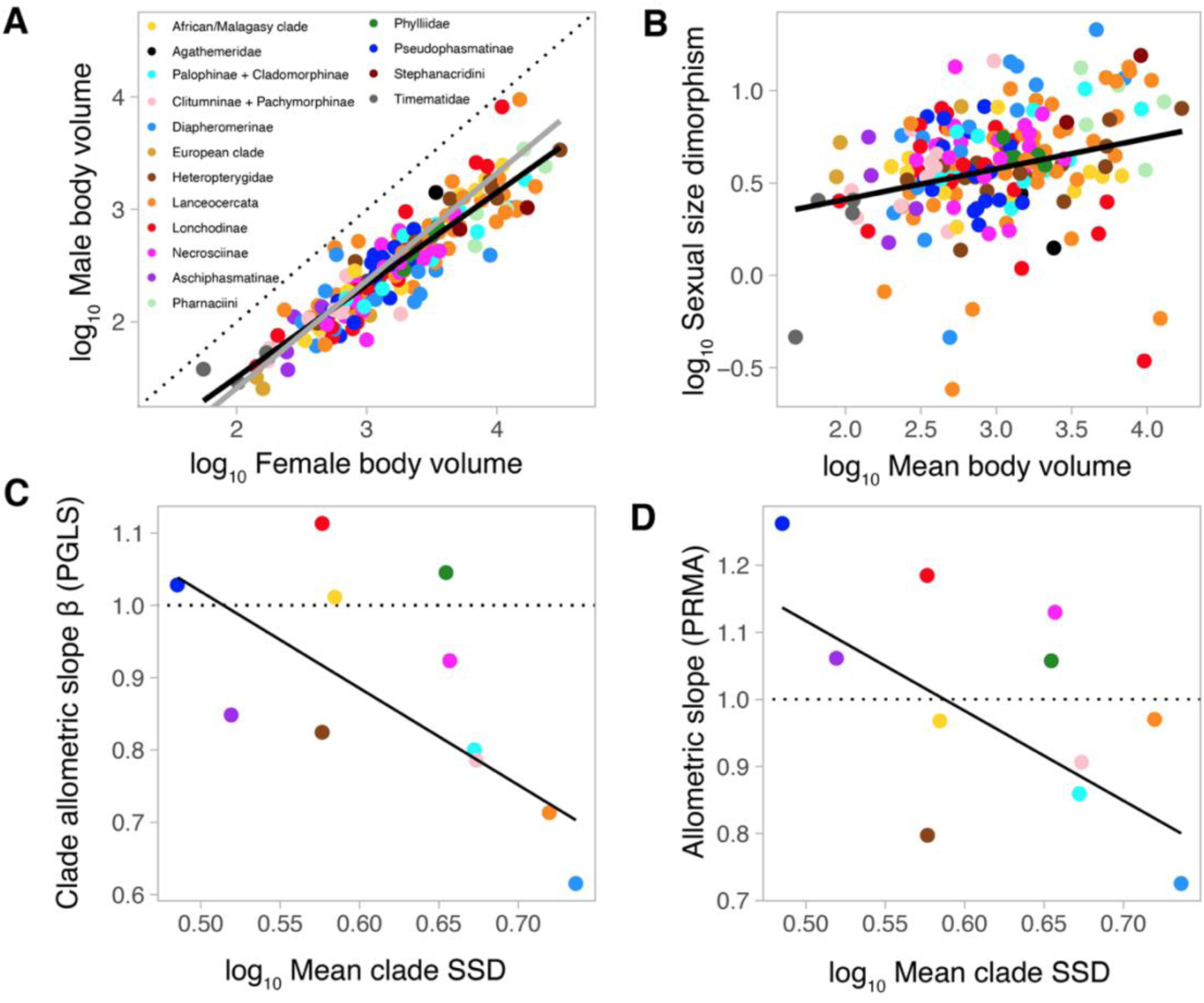
Sexual size allometry in stick insects (Phasmatodea). **A.** Log_10_ male body volume versus log_10_ female body volume (mm^3^) with PGLS (black) and PRMA (gray) regressions shown. Colors correspond to the main phasmid clades. **B.** Log_10_ Sexual size dimorphism (SSD) versus log_10_ mean body volume of males and females. **C-D**. Allometric slopes β, calculated from clade-specific PGLS (**C**) or PRMA (**D**), versus phylogenetically controlled mean SSD by major phasmid clade (see color legend in **A**.). The dashed lines correspond to isometry (1:1 relationship). Associated clade-specific data can be found in Table S1.

A PGLS regression between log_10_ male body volume and log_10_ female body volume across all phasmid species showed an allometric slope 𝛽 of 0.82 that significantly departed from 1 (Figure 2A, 𝜆= 0.79, 95% confidence interval (CI): 0.76–0.89), thereby supporting the inverse of Rensch’s rule. However, the corresponding PRMA exhibited a 𝛽 of 0.96, which was not significantly different from isometry (Figure 2A, 𝜆= 0.96, 95% CI: 0.88–1.03). Finally, we found a positive relationship between SSD and the mean species body volume (Figure 2B, PGLS: 𝜆=0.62, slope=0.16 ± 0.05, F_1,194_ =10.78, p=0.001), again going against the definition of Rensch’s rule.

In accordance with the predictions of quantitative genetic theory (Zeng 1988; De Lisle and Rowe 2013), we found a significant negative relationship between clade-specific phylogenetically corrected mean SSD values and the associated 𝛽 values calculated with the PRMA (F_1,9_= 5.16 , p=0.049, Figure 2C) or PGLS (F_1,9_= 7.37, p=0.024, Figure 2D) methods.

This suggests that, historically, the evolution of female-biased SSD in stick insects has been primarily driven by positive directional selection on female size.

### Proximate causes of SSD

SSD was positively correlated with sex difference in development duration and sex difference in relative growth rate (Table 1, Figure 3). The multiple-predictor PGLS further revealed that scaled sex differences in growth rate and in duration of development had similar effect sizes on the extent of SSD (Table 1), thus indicating that females are larger than males in phasmids because they develop faster and for longer, with equal importance.

**Figure 3:**
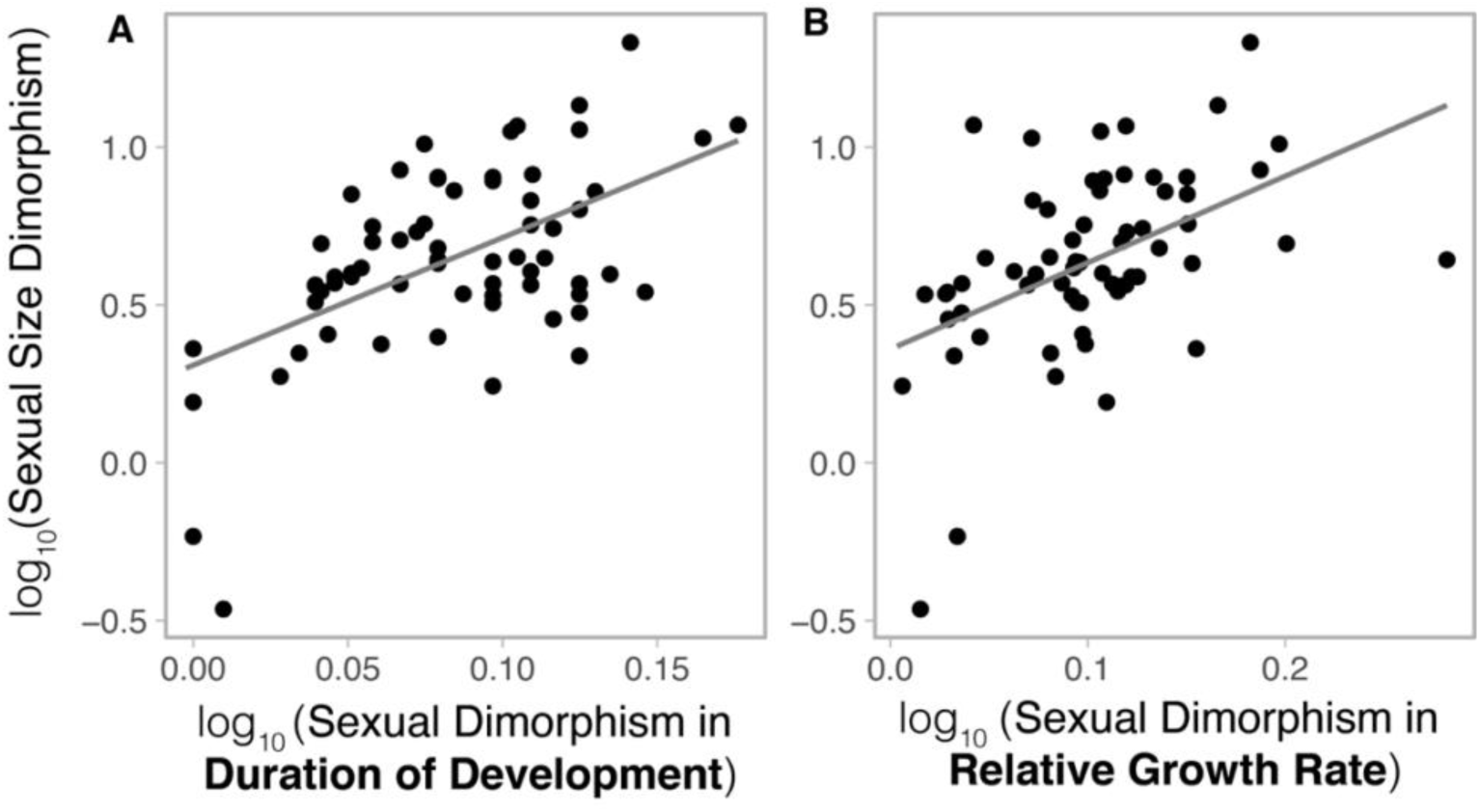
Sex differences in duration of postembryonic development (SDDD) and relative growth rate (SDRGR) explain sexual size dimorphism (SSD). Log_10_ (SSD) versus log_10_ (SDDD) (**A**) and log_10_ (SDRGR) (**B**), with PGLS regressions shown. Details of the associated statistical tests can be found in Table 1.

**Table 1:** Sexual size dimorphism is caused by both sex differences in postembryonic development duration and relative growth rate. The table presents results of three phylogenetic generalized least squares models (PGLS) --and corresponding type III ANOVA-- including sexual size dimorphism (SSD) as the response variable (log10-transformed) and sexual dimorphism in duration of development (SDDD) and/or in relative growth rate (SDRGR). SDTD and SDRGR were log10-transformed, centered on zero and scaled to unit variance. Bold characters highlight significant effects (p<0.05).

| PGLS | Pagel's $\lambda$ | Response | Explanatory variable | Effect size (± SE) | F<sub>df1, df2</sub> | P-value |
| --- | --- | --- | --- | --- | --- | --- |
| Single-predictor | 0.30 | $\log_{10}$ (SSD) | **$\log_{10}$ (SDDD)** | **$0.16 \pm 0.03$** | **30.0**<sub>1,65</sub> | **<0.0001** |
| Single-predictor | 0.02 | $\log_{10}$ (SSD) | **$\log_{10}$ (SDRGR)** | **$0.14 \pm 0.03$** | **18.6**<sub>1,62</sub> | **0.0001** |
| Multiple-predictor | 0.18 | $\log_{10}$ (SSD) | **$\log_{10}$ (SDDD)** | **$0.20 \pm 0.02$** | **84.1**<sub>1,60</sub> | **<0.0001** |
|  |  |  | **$\log_{10}$ (SDRGR)** | **$0.18 \pm 0.02$** | **77.4**<sub>1,60</sub> | **<0.0001** |
| | | | interaction | $-0.02 \pm 0.02$ | 0.51<sub>1,60</sub> | 0.48 |

### Selective drivers of variation in SSD

Single and multiple predictor PGLS models showed that female reproductive output (fecundity) was positively related to both female and male size, but not to SSD (Table S2, Figure 4A-D), thereby rejecting hypothesis I. Female fecundity alone explained 67% of the variation in female volume and 51% of the variation in male volume, but 0% of the interspecific variation in SSD (Table S2), after accounting for phylogenetic relatedness. This result seems contradictory with the previous finding that female-biased SSD in phasmids resulted from historically stronger positive selection on females. Therefore, it suggests that fecundity selection and overall positive selection on female size has driven an average female-biased SSD in phasmids but that variation in SSD around that average cannot be explained by variation in female fecundity, suggesting that other factors may be at play.

**Figure 4:**
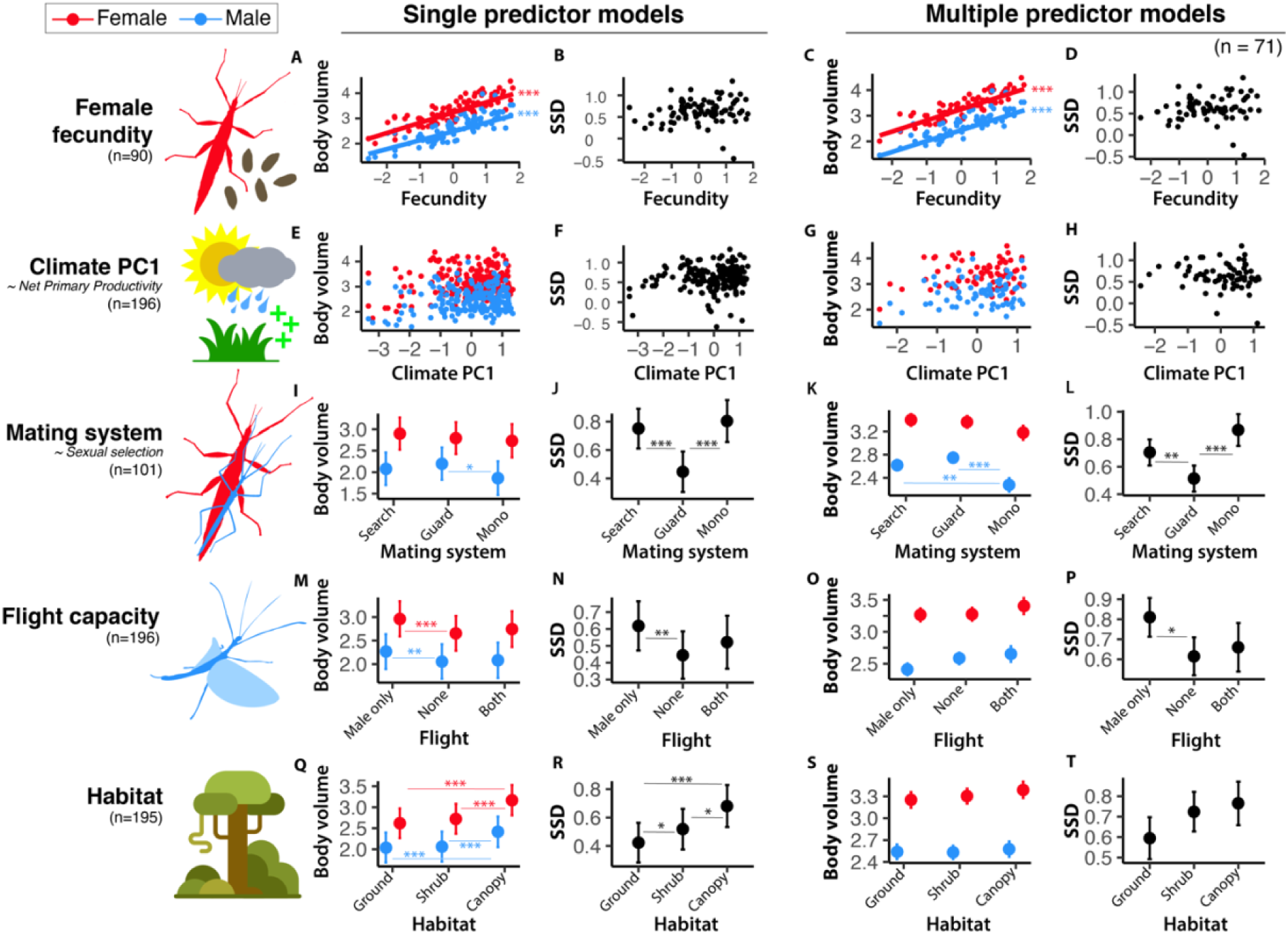
Ultimate predictors of male and female body size and sexual size dimorphism (SSD). Results of single predictor (left columns, sample size indicated below each predictor) and multiple predictor (right columns, n=71 species) PGLS models are presented. Log_10_ male body volume (blue, mm^3^), log_10_ female body volume (red, mm^3^) and log_10_ SSD (black) are shown versus log_10_ female fecundity (i.e., lifetime reproductive output) (**A-D**), climate PC1 (**E-H**), mating system (**I-L**), flight dimorphism (**M-P**) and habitat (**Q-T**). PGLS regressions with significant slopes are shown for continuous predictors (𝛽 ≠ 0, p <0.05) (**A-H**). Estimated marginal means (± standard error) for each group are shown for categorical predictors (**I-T**). Significant pairwise group differences (assessed using the Holm correction) are shown. Asterisks indicate significant slopes/group differences (*: p <0.05; **: p<0.01, ***: p<0.001). Corresponding statistical analyses can be found in Table S2.

Both model types also showed that mating system significantly affected male but not female size, thus affecting SSD (Table S2, Figure 4I-L). These findings supported hypothesis II and were consistent with our predictions: guarding males are larger than non-guarding searching males and monogamous males. Mating system alone explained 0% of the variation in female volume and 5% of the variation in male volume, but 29% of the variation in SSD (Table S2), after accounting for phylogenetic relatedness.

We did not find any correlation between climate and male or female body volume or SSD (Table S2, Figure 4E-H), suggesting that ecological selection resulting from limited food availability or growing season length did not account for species differences in SSD.

Single predictor models revealed that males and females were significantly larger in species where only males are able to fly (Table S2, Figure 4M), which went against our prediction that flight-capable males would be smaller than flightless ones. Nevertheless, this effect was stronger in females than males resulting in a significantly higher SSD in species with flight capable males and flightless females (Table S2, Figure 4N), which was consistent with our original predictions. Flight dimorphism alone explained 9% of the variation in female volume, 4% of the variation in male volume, and 5% of the variation in SSD (Table S2), after accounting for phylogenetic relatedness. Our multiple predictor models similarly revealed that species with flying males and flightless females were more dimorphic than species with both sexes being flightless (Table S2, Figure 4P). However, they showed that this difference in SSD was mainly caused by flight-capable males being smaller in species with only males able to fly and not by females being larger (Table S2, Figure 4O), thus agreeing with our predictions. Species with both sexes able to fly, for which we predicted a reduced size of males and females and a low SSD, did not differ from other categories in any of the models probably owing to a reduced sample size (n= 18 species, only including 9 with sufficient data to be included in the multiple predictor models).

Single predictor models showed that species dwelling in higher vegetation strata had a higher SSD, owing to a stronger effect on female than male size (Table S2, Figure 4Q-R). Habitat alone explained 16% of the variation in female volume, 8% of the variation in male volume and 7% of the variation in SSD (Table S2), after accounting for phylogenetic relatedness. In contrast, multiple predictor models did not show any significant effect of habitat on male or female volume or SSD. This may be partly explained by the correlation between habitat and flight dimorphism as volant forms (especially in males) are more commonly found higher up in the vegetation.

In the multiple predictor models including female volume, male volume and SSD as response variables, the variance explained by both phylogeny and predictors was higher (R^2^ (SSD)= 0.43, R^2^ (female volume)= 0.82, R^2^ (SSD)= 0.75) than that explained by the predictors alone (R^2^ (SSD)= 0.40, R^2^ (female volume)= 0.76, R^2^ (SSD)= 0.71, Table S2). But most of the variation was explained by fixed effects alone and not by phylogenetic relationships per se. This suggests that body size is quite phylogenetically labile.

### Selective drivers of variation in SShD

While single predictor models showed significant effects of mating system, flight dimorphism and habitat on male and female body elongation (PC1)(Table S3), multiple predictor models, accounting for the intercorrelation between predictors, only found a significant effect of mating system on PC1 in males and consequently on SShD_elongation_ (Figure 5A-J, Table S3). Guarding males, whether they only temporarily guard females or guard them for their lifetime, tended to be stockier and less elongated than purely searching males, resulting in the most dimorphic overall body silhouettes being found in species with searching males and sedentary females.

**Figure 5:**
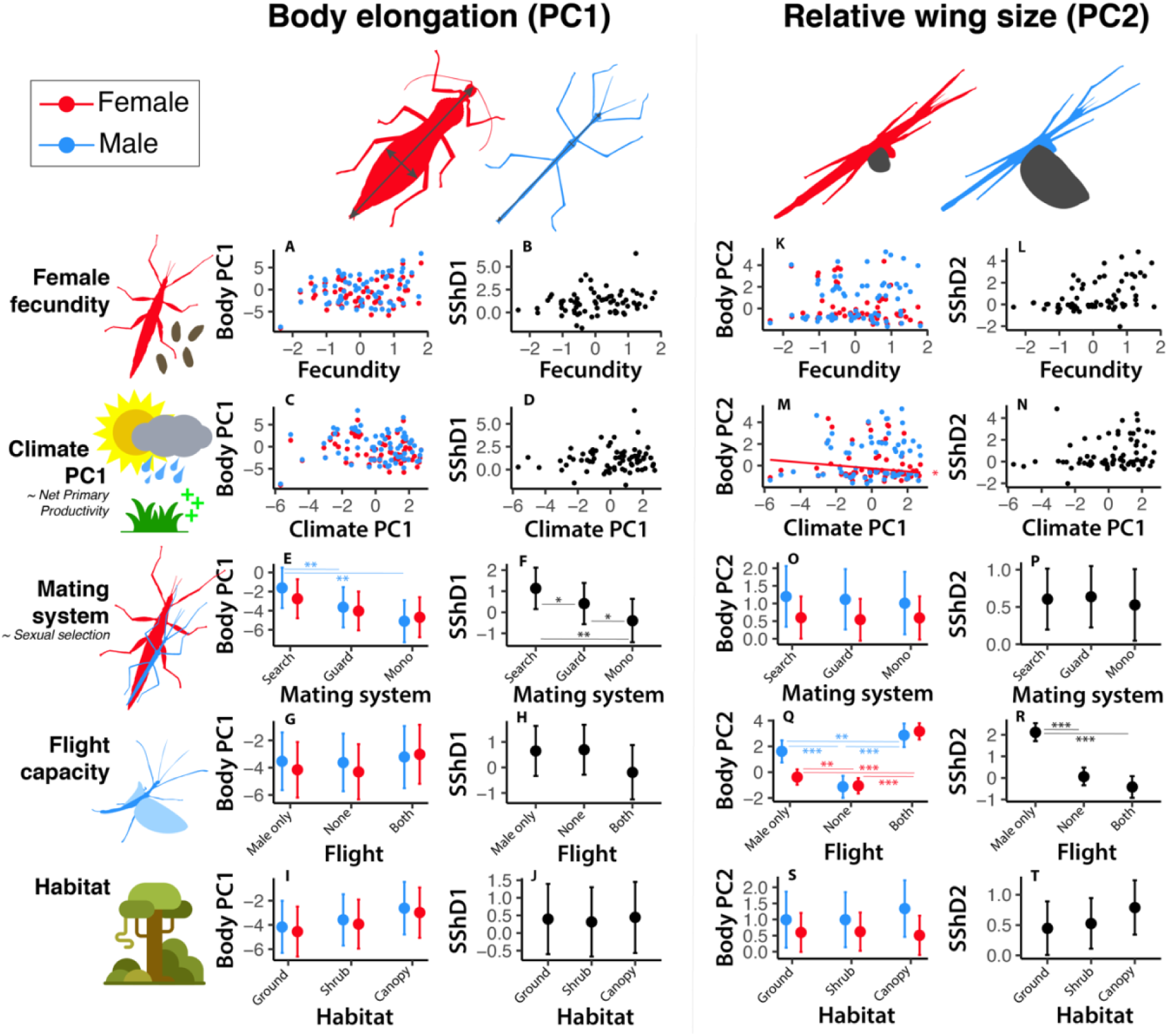
Ultimate predictors of male and female body elongation (PC1) and relative wing size (PC2), and associated sexual shape dimorphism (SShD _elongation_ and SShD _wings_). Results of multiple predictor PGLS models (n= 69 species) are presented. Male (blue) and female (red) body PC1, SShD _elongation_ (**A-J**), male (blue) and female (red) body PC2 and SShD _wings_ (**K-T**) are shown versus log_10_ female fecundity (i.e., lifetime reproductive output) (**A-B, K-L**), Climate PC1 (**C-D, M-N**), Mating system (**E-F, E-P**), Fight dimorphism (**G-H, Q-R**) and habitat (**I-J, S-T**). Significant slope coefficients are shown for continuous predictors (𝛽 ≠ 0, p <0.05) (**A-D, K-N**). Estimated marginal means (± standard error) for each group are shown for categorical predictors (**E-J, O-T**). Significant pairwise group differences (assessed using the Holm correction) are shown. Asterisks indicate significant slopes/group differences (*: p <0.05; **: p<0.01, ***: p<0.001). Corresponding statistical analyses can be found in Table S3-S4.

In the multiple predictor models including female PC1, male PC1 and SSD_elongation_ as response variables, the variance explained by both phylogeny and predictors was higher (R^2^ (SSD)= 0.37, R^2^ (female volume)= 0.55, R^2^ (SSD)= 0.59) than that explained by the predictors alone (R^2^ (SSD)= 0.09, R^2^ (female volume)= 0.45, R^2^ (SSD)= 0.34, Table S2). Here, the part of the variance explained by phylogeny was higher than for body size suggesting that body elongation is more phylogenetically conserved.

Similarly, single predictor models suggested a significant effect of mating system, flight dimorphism and habitat on male and female relative wing size (PC2)(Table S4), but multiple predictor models only found a significant effect of flight dimorphism (Figure 5K-T, Table S4). As expected, flying males and females had relatively larger wings than non-flying ones. Consequently, dimorphism in wing size was much greater in species where only males can fly. Finally, one of our multiple predictor models found a significant effect of climate on female PC2. It suggested that, after accounting for the other factors, females have relatively shorter wings in more tropical macrohabitats (i.e., with higher net primary productivity).

In the multiple predictor models including female PC2, male PC2 and SSD_wings_ as response variables, the variance explained by both phylogeny and predictors was sometimes lower (R^2^_full_ (SSD)= 0.68, R^2^_full_ (female volume)= 0.86, R^2^_full_ (SSD)= 0.82) than that explained by the predictors alone (R^2^_fixed_ (SSD)= 0.63, R^2^_fixed_ (female volume)= 0.93, R^2^_fixed_ (SSD)= 0.87, Table S2). This clearly suggests that relative wing size is not phylogenetically conserved and varies substantially between sister taxa.

## Discussion

The stick and leaf insects (Phasmatodea) display breathtaking interspecific variability in overall body size and in the extent and nature of sexual dimorphism. Here we tested three alternative hypotheses for the evolution of species differences in sexual dimorphism in both body size (SSD) and shape (SShD): (I) fecundity selection favoring larger and thicker females, (II) sexual selection favoring larger and stockier males, and (III) ecological variables differentially affecting male and female body size and shape. We show that positive selection on female size (likely stemming from fecundity selection) has been the primary historical driver of female-biased SSD in phasmids. However, in contrast with other studied female-biased taxa (e.g., amphibians (Monroe et al. 2015), fishes (Horne et al. 2020)), fecundity selection does not appear to have driven contemporary species differences in sexual dimorphism in these insects (rejecting hypothesis I). Instead, interspecific patterns of SSD and SShD likely resulted from selection acting on the males, either sexual selection (species differences in mating system, supporting hypothesis II) or ecological selection related to flight capability (supporting hypothesis III). Consequently we propose that (1) positive selection on female size has been the main historical driver of female-biased SSD in stick insects and that fecundity selection consistently favors larger female sizes across taxa but that (2) contemporary variation around the average female-biased SSD is mainly caused by selection acting on male size, whose magnitude and direction may be more variable across species.

Directional selection for larger female size is most often expected to result from the positive correlation between body size and fecundity (Fairbairn 1997; Pincheira-Donoso and Hunt 2017). Accordingly, we found a strong and positive correlation between female lifetime reproductive output and female size across stick insects (Figure 4A). However, and contrary to our expectations, variation in female fecundity was not correlated with variation in SSD (Figure 4B), in contrast with other female-biased systems including fishes (Horne et al. 2020) and amphibians (Monroe et al. 2015). While we cannot predict the magnitude of SSD based on female fecundity, it is still likely a major factor maintaining large female sizes and driving SSD. Indeed, our data from 10 major phasmid clades showed that the magnitude of SSD of a clade increases as females become more size-variant than males (i.e., lower 𝛽, Figure 2C-D). Consistent with predictions from quantitative genetic theory (Zeng 1988; De Lisle and Rowe 2013), this pattern suggests that directional selection acting on female size played a major role in generating hypoallometry (β < 1) and female-biased SSD in stick insects. In other words, this implies that positive selection on female size, rather than negative selection on male size, may have been the prominent driver of species size and female-biased SSD in phasmids. Therefore, the intensity of fecundity selection may not vary substantially across species and, while strong fecundity selection may have historically driven the evolution of much larger females, deviations observed today from the average female-biased SSD end up being mostly explained by variation in selective pressures acting on male size. Indeed, we start seeing males and females of comparable sizes (but never reaching male-biased SSD or monomorphism) only when pressures favoring larger male sizes become relatively intense, as exemplified by the weapon-bearing thorny devil stick insects. Unlike most other stick insect species, the relatively large male thorny devil stick insects (*Eurycantha* spp.) harbor extremely enlarged hindlegs that they use in violent combats with rival males over females or territories (Boisseau et al. 2020; Boisseau and Emlen 2025). Here, we found that female *E. calcarata* are only 1.34 times larger than males, relative to a phylogenetically-corrected average of 3.96 across all phasmids. We however recognize that the intensity of fecundity selection in a given taxa would be better quantified as the taxon-specific slope of the scaling relationship between female reproductive output and body size (Horne et al. 2020). Ultimately, our results suggest that although selection acting on male size best predicts variation in SSD in phasmids, fecundity selection acting on female size was likely the primary driver of the ubiquitous female-biased SSD seen in the group.

Our analyses pointed out to mating system (i.e., sexual selection intensity) and flight capacity as being the best predictors of variation in the extent of SSD and SShD across phasmids. In species where males do not typically stay with a female for extended periods of time, male reproductive success is likely determined by their searching and locomotor performance, which are predicted to be enhanced with smaller sizes (Kelly et al. 2008; Herberstein et al. 2017; Boisseau et al. 2020, 2022; Kelly 2020). Consistently, males of such species are relatively smaller and slenderer (Figure 4I-L & 5E-F), even after accounting for their flight capacity. This suggests that even walking males may benefit from being smaller, maybe owing to a higher substrate attachment performance (Boisseau et al. 2022) or a lower energy need which would translate in a lower need to allocate time to foraging relative to mate searching (i.e., “Ghiselin–Reiss small-male hypothesis” (Ghiselin 1974; Reiss 1989; Blanckenhorn et al. 1995)). In contrast, species in which males search for and subsequently guard females have relatively larger and stockier males (Figure 4I-L & 5E-F). In such species, the post-copulatory mate guarding behavior of males suggests a high risk of sperm competition and a higher likelihood of one-on-one male encounters and physical contests (Sivinski 1979; Boisseau et al. 2020; Wilner et al. 2025). In the prairie walkingstick (*Diapheromera velii*, Diapheromerinae) and other similar mate guarding systems, the largest and strongest males are almost invariably more successful at interrupting and displacing other guarding males and, when guarding themselves, are better able to resist takeovers (Sivinski 1979; Borgia 1980; Bel-Venner and Venner 2006; Knox and Scott 2006). Although male reproductive success is likely dependent on guarding ability in these species, in most cases, those males still extensively search for scattered females (Sivinski 1977; Myers et al. 2015), and consequently their body morphology likely reflects the balance between selective pressures stemming from both searching and fighting (Kelly 2014). The relative importance of these two factors may vary between species and also temporarily and spatially with the operational sex ratio and the relative densities of males and females (Schöfl and Taborsky 2002; Kelly 2015; Myers et al. 2015; Simmons et al. 2020). A more detailed quantification of the relative importance of searching and guarding would be desirable to better quantify variation in sexual selection intensity. Finally, monogamous males which stay on the back of a female for most of their lifetime exhibit much smaller sizes but a similar body shape compared to their respective females (Figure 4I-L & 5E-F). By staying in copula for very long periods of time, these males’ body essentially acts as a sperm plug insuring exclusive paternity of the female’s offspring. Thus, the reproductive interests of the two sexes should align and therefore selection should favor male traits —e.g., small body sizes— that help reduce the handicap of carrying its mate for the female. Dwarfed males may also benefit from lessening the disruption of camouflage while riding females and from keeping their maintenance cost to a minimum. Direct physical combat from intruding solitary rivals may be infrequent as solitary males approaching already coupled pairs have been observed to simply withdraw in such systems (e.g., *Anisomorpha buprestoides,* Pseudophasmatinae (Gunning 1987)), suggesting a high owner’s advantage and a low probability of takeovers potentially relieving selection for physical strength and larger sizes.

Differences in flight abilities between sexes was also a significant predictor of the magnitude of SSD and, unsurprisingly, SShD_wings_. Indeed, after accounting for confounding factors such as mating system, flight appeared to favor smaller body sizes and relatively larger wings, leading to a lower wing loading. Wing loading –i.e., body mass divided by wing area–is often a good predictor of flight speed, wing beat frequency and therefore flight cost. Flapping animals with a high wing loading must fly fast to generate enough lift to stay in the air while animals with a low wing loading can fly slowly and inexpensively and benefit from an increased maneuverability (Biewener and Patek 2018; Le Roy et al. 2019). Stick and leaf insects are slow fliers for which a higher wing loading translate into a reduced climbing ability and maneuverability in flight, and therefore a reduced flight performance and capability (Boisseau et al. 2022; Zeng et al. 2023). Wings in stick insects can also be used in other contexts unrelated to flight including parachuting and predator deterrence through threatening displays of showy colorations (Zeng et al. 2020). We could have expected flightless but winged species living in high vegetation layers to harbor relatively larger wings to slow down hypothetical falls, even if they cannot actively fly. But no other factor except flight capacity affected relative wing size.

Our data shows that female size is overall more variable than male size (𝛽 < 1) across Phasmatodea and that SSD increases with species size, thereby supporting the inverse of Rensch’s rule. In addition, clade-specific allometries were never significantly hyperallometric and therefore never followed Rensch’s rule (Table S1). These results are generally consistent with other groups with female-biased SSD, such as various insects (Blanckenhorn et al. 2007b) (including a previous study on stick insects using body length as body size metric (Sivinski 1979; Sivinski and Dodson 1992)), birds (Webb and Freckleton 2007) and fishes (Horne et al. 2020).

SSD can arise from sex differences in post-embryonic development time and/or growth rate. The relative contributions of each of these proximate mechanisms has already received some attention, especially in insects (Blanckenhorn et al. 2007a; Teder 2014; Teder et al. 2021) and fishes (Horne et al. 2020). We found that, in stick insects, species with the greatest relative difference in development time and in relative growth rate between sexes were the most size dimorphic. Females developed faster and for longer than males, both mechanisms conjointly generating female-biased SSD with nearly equal importance. Protandry –i.e., males maturing earlier than females– is ubiquitous in stick insects with only 3 out of 67 species with development time data exhibiting negligible differences in development duration. It is also the most common pattern of sex difference in development time in insects, and its evolution appears to be mainly driven as a by-product of the evolution of SSD (Teder et al. 2021). Nevertheless, it may still be advantageous for male stick insects to mature earlier. In taxa with discrete and nonoverlapping generations in seasonal environments (e.g., *Timema*, *Diapheromera*, *Bacillus*), males maturing early may have a mating advantage (Wiklund and Fagerström 1977; Allen et al. 2011; Teder et al. 2021). For instance, early male maturation may be beneficial in systems where males guard pre-mature females (e.g., *Orthomeria kangi*, Aschiphasmatinae (Vallotto et al. 2016)) or where males pair with females for a very long time and, as discussed above, are unlikely to be overthrown by rivals once anchored (e.g., *Anisomorpha buprestoides*, Pseudophasmatinae (Gunning 1987)). But such advantage should disappear in species with overlapping generations where mature females are continuously available, as is the case for a lot of rainforest species. It should finally be noted that stick insects are generally sedentary and, despite some remarkable adaptations to dispersal (notably in the eggs (O’Hanlon et al. 2020; Boisseau and Woods 2024)), may be at risk of inbreeding depression. Males maturing earlier, and sometimes dying before their sisters even reach maturity, may avoid this risk (Wiklund and Fagerström 1977).

## Conclusion

To date, the majority of large-scale comparative analyses investigating the extent to which SSD is driven by selection for increased size in males versus females, have consistently found correlation between the extent of SSD and the intensity of sexual selection acting on male size (Serrano-Meneses and Székely 2006; Lislevand et al. 2009; Monroe et al. 2015; Horne et al. 2020), but, interestingly, have rarely offered support for the often-evoked fecundity selection hypothesis. However, our study exemplifies that, while variation in SSD may not be predictable based only on variation in female fecundity, it does not mean it is irrelevant to the evolution of SSD. Allometric patterns of SSD in stick insects suggest that positive selection on female size has been the most important historical driver of female-biased SSD in the group. Thus it suggests that fecundity may consistently favor larger female sizes across taxa but that variation around the average SSD is mainly caused by selection acting on male size, whose magnitude and direction may be more variable across species. By combining descriptions of allometric patterns of SSD and analyses of the effects of proximate and ultimate predictors on SSD and SShD, we hope our study can serve as a methodological template for future studies investigating the diversity of sexual dimorphism in other groups.

---

**Figure S1:**
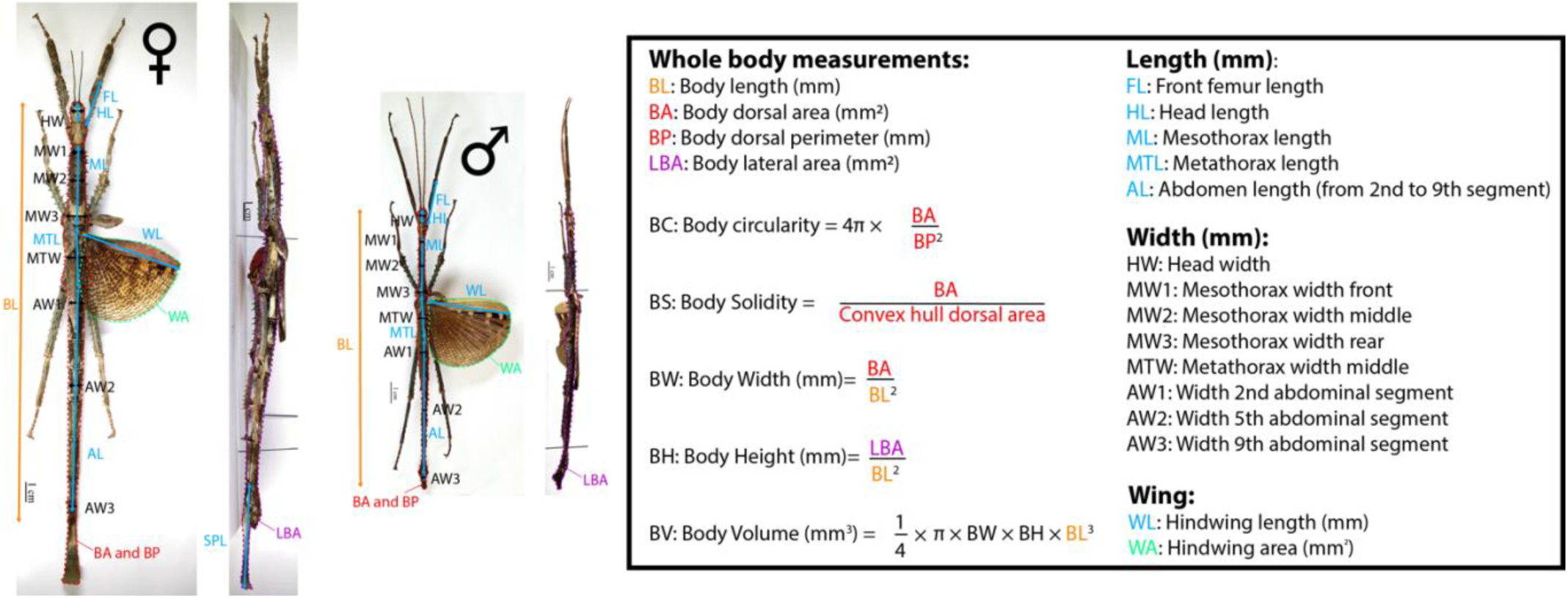
Measurements of phasmid specimens. **Left:** Adult female *Achrioptera punctipes cliquennoisi* Hennemann & Conle 2004 (Specimen MNHN-EO-PHAS127, project RECOLNAT (ANR-11-INBS-0004), photographs by Marion Depraetere, 2015, CC-BY-NC-ND) in dorsal and lateral view. **Middle:** Adult male *Achrioptera punctipes cliquennoisi* Hennemann & Conle 2004 (Specimen MNHN-EO-PHAS126, project RECOLNAT (ANR-11-INBS-0004), photographs by Marion Depraetere, 2015, CC-BY-NC-ND) in dorsal and lateral view. Body perimeter was only used to calculate body circularity and was not included in the principal component analysis described in figure S2. Orange corresponds to body length (measured in absolute). Light blue corresponds to body part length measurements (measured relative to body length). Black corresponds to body part width measurements (measured relative to body length). Red corresponds to body dorsal perimeter and area (measured relative to body length). Purple corresponds to body lateral area (measured relative to body length). Green corresponds to wing area (measured relative to body length).

**Figure S2:**
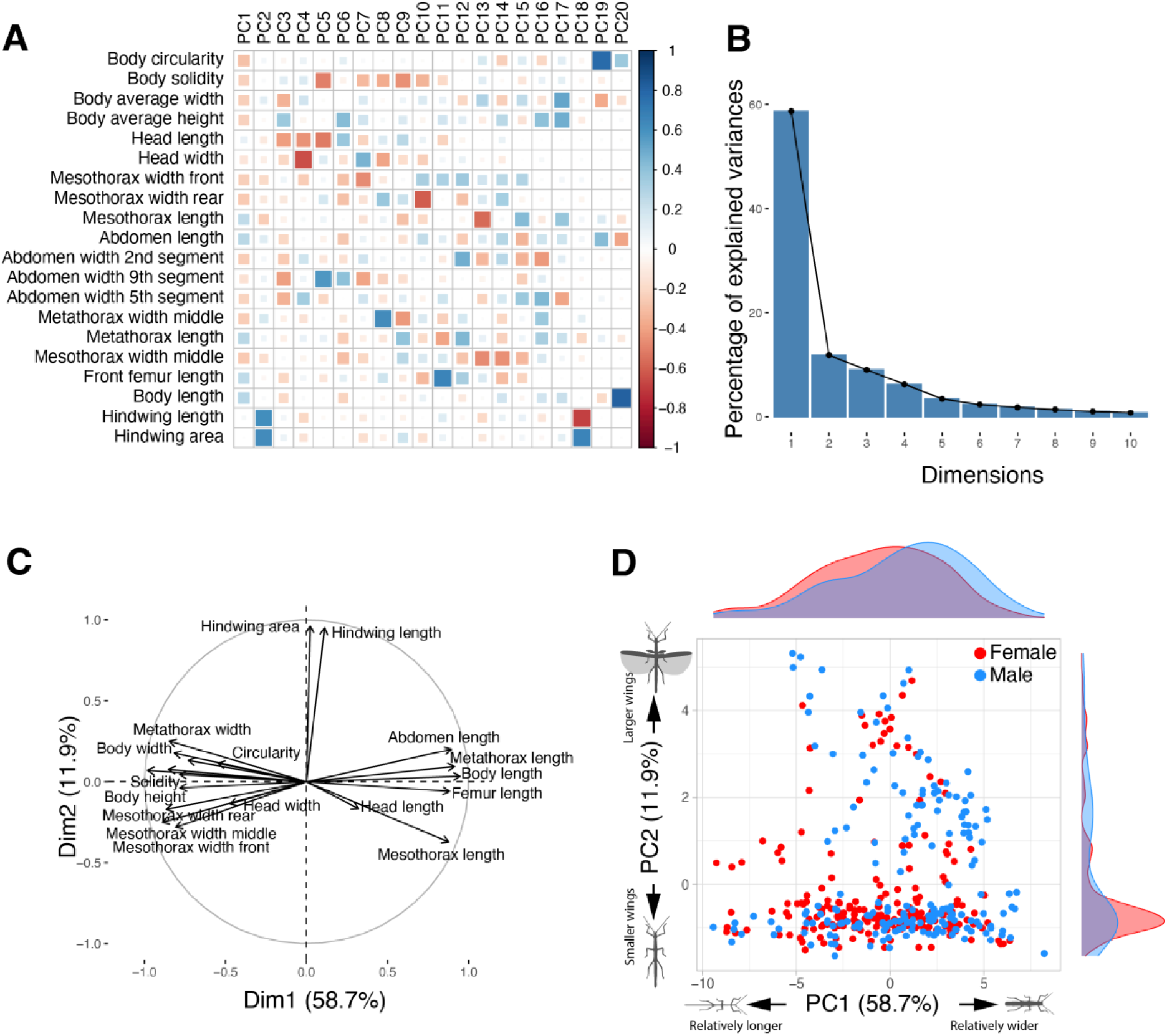
Principal component analysis (PCA) summarizing body shape of males and females. **A**: Correlation plot showing the extent and direction of the correlations between each principal component (PC) and the body shape variables included (i.e., residuals of PGLS regressions against body volume). **B**: Proportion of the total variance explained by each PC. **C**: PCA loading plot showing how strongly each characteristic influences PC1 and PC2. **D**: Morphospace showing PC2 against PC1 for males (blue) and females (red) of each species included.

**Figure S3:**
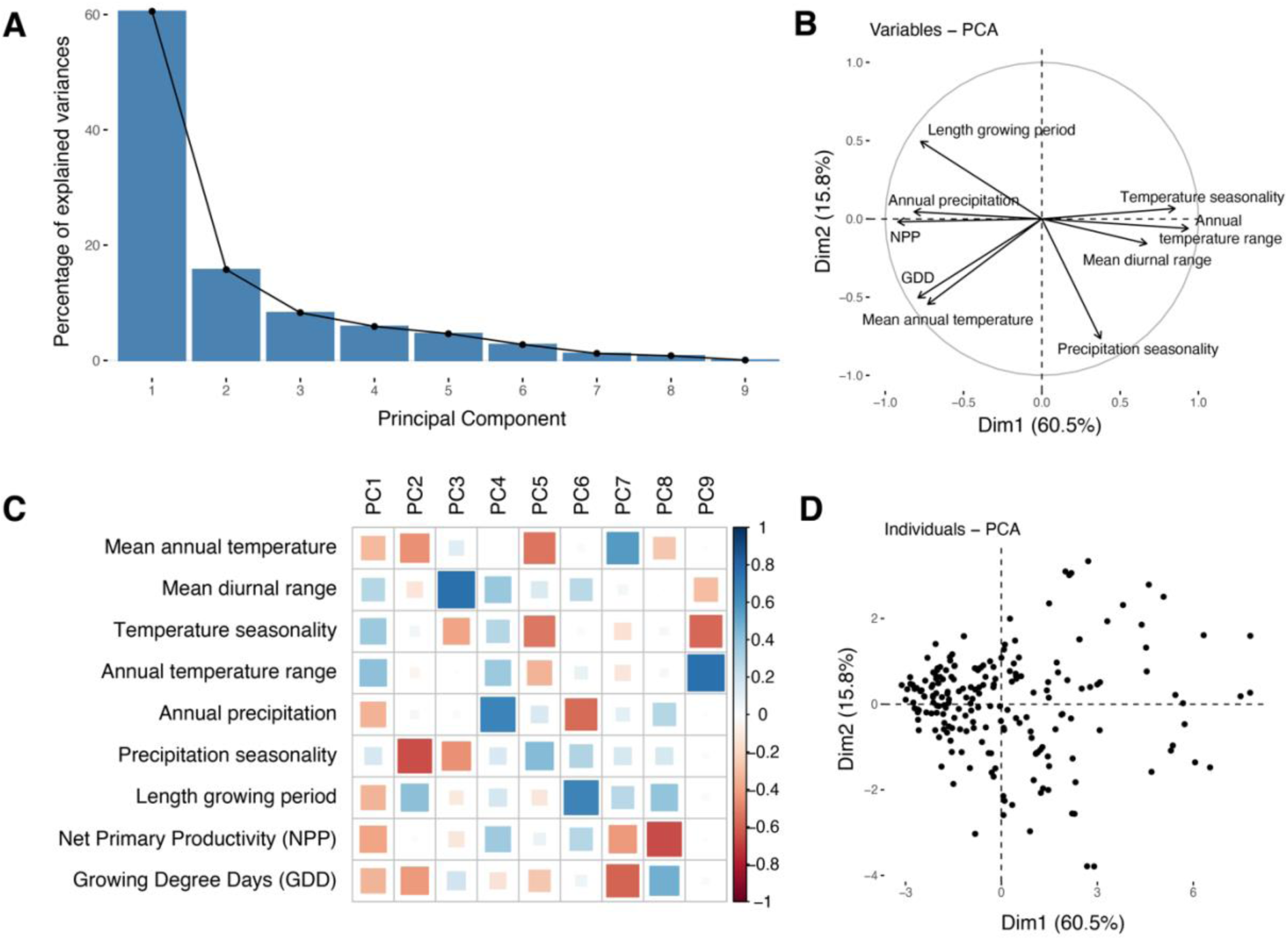
Principal component analysis (PCA) summarizing climatic conditions encountered by phasmid species. **A**: Proportion of the total variance explained by each principal component (PC). **B**: PCA loading plot showing how strongly each climatic variable influences PC1 and PC2. **C**: Correlation plot showing the extent and direction of the correlations between each principal component (PC) and the climatic variables included. **D**: PC2 against PC1 for each species included.

**Figure S4.**
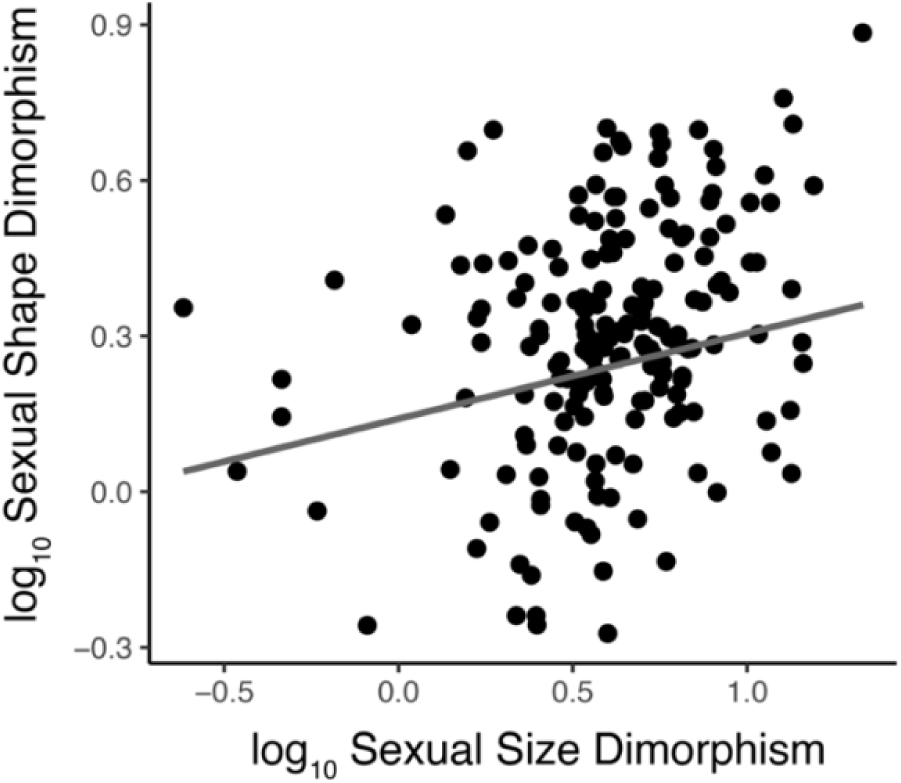
Phylogenetically controlled correlation between overall SShD and SSD (PGLS regression: 𝜆 = 0.59, F_1,184_ = 9.82, p = 0.002, r^2^ = 0.05).

**Table S1:** Clade-specific phylogenetically corrected mean SSD and allometric slope (calculated using a pRMA or PGLS) between log_10_ male and female body volume, and associated 95% confidence intervals.

| Clade | N<sub>species</sub> | RMA slope | RMA 95% CI | PGLS slope | PGLS 95% CI | Mean SSD |
| --- | --- | --- | --- | --- | --- | --- |
| Aschiphasmatinae | 6 | 1.06 | 0.33-1.80 | 0.85 | 0.22-1.47 | 3.31 |
| Diapheromerinae | 14 | **0.73** | 0.45-1.00 | **0.62** | 0.37-0.86 | 5.45 |
| Pseudophasmatinae + Agathemeridae | 19 | 1.26 | 0.99-1.54 | 1.03 | 0.76-1.30 | 3.06 |
| Heteropteryginae | 17 | **0.80** | 0.66-0.94 | **0.82** | 0.72-0.93 | 3.77 |
| African/Malagasy/<br>European clade | 15 | 0.97 | 0.77-1.17 | 1.01 | 0.88-1.15 | 3.80 |
| Phylliinae | 6 | 1.06 | 0.63-1.49 | 1.05 | 0.89-1.20 | 4.51 |
| Necrosiinae | 18 | 1.13 | 0.75-1.51 | 0.92 | 0.61-1.24 | 4.54 |
| Lonchodinae | 23 | 1.18 | 0.94-1.43 | 1.11 | 0.93-1.30 | 3.77 |
| Palophinae + Cladomorphinae | 12 | 0.86 | 0.62-1.09 | **0.80** | 0.61 - 0.99 | 4.70 |
| Pharnaciini + Clitumninae + Pachymorphinae | 16 | 0.91 | 0.61-1.20 | **0.79** | 0.65-0.92 | 4.72 |
| Stephanacridini + Lanceocercata | 46 | 0.97 | 0.80-1.14 | **0.71** | 0.59-0.83 | 5.24 |

**Table S2:** Ultimate predictors of variation in female and male volume and SSD. The table presents results of single and multiple predictor PGLS models. The most likely value of Pagel’s lambda (phylogenetic signal) is presented along with ANOVA outputs (using marginal sum of squares obtained by deleting a term from the model at a time) and either estimated effect sizes or post-hoc pairwise comparisons between estimated marginal means using the Holm method to account for multiple testing, respectively for continuous or categorical explanatory variables. The proportions of the variance explained by the full model R^2^_full_ (phylogeny + fixed effects) and explained by fixed effects alone R^2^_fixed_ (after accounting for phylogeny) are also reported.

| # predictors | Response | Predictor | $\lambda$ | $R^2_{full}$ | $R^2_{fixed}$ | $F_{df1,df2}$ | P | Effect size ± Standard Error (continuous)<br>OR<br>Post-hoc pairwise tests (Holm) (categorical) |
| --- | --- | --- | --- | --- | --- | --- | --- | --- |
| 1 | $\log_{10}$ female volume | **$\log_{10}$ Female reproductive output** | 0.42 | 0.77 | 0.67 | $F_{1,88} = 185.8$ | <.0001 | **0.41 ± 0.03** |
| | $\log_{10}$ male volume | **$\log_{10}$ Female reproductive output** | 0.57 | 0.68 | 0.51 | $F_{1,88} = 94.9$ | <.0001 | **0.35 ± 0.04** |
| | $\log_{10}$ SSD | $\log_{10}$ Female reproductive output | 0 | 0 | 0 | $F_{1,88} = 2.31$ | 0.13 | 0.043 ± 0.03 |
| 1 | $\log_{10}$ female volume | Climate PC1 | 0.99 | 0.54 | 0.01 | $F_{1,194} = 3.38$ | 0.07 | 0.065 ± 0.036 |
| | $\log_{10}$ male volume | Climate PC1 | 0.99 | 0.49 | 0 | $F_{1,194} = 1.17$ | 0.28 | 0.037 ± 0.034 |
| | $\log_{10}$ SSD | Climate PC1 | 0.69 | 0.21 | 0 | $F_{1,194} = 0.85$ | 0.36 | 0.023 ± 0.025 |
| 1 | $\log_{10}$ female volume | Mating system | 0.95 | 0.48 | 0 | $F_{2,98} = 1.13$ | 0.33 | Search – Guard = 0.11 ± 0.08, p = 0.62<br>Search – Mono = 0.17 ± 0.14, p = 0.62<br>Guard – Mono = 0.06 ± 0.14, p = 0.65 |
| | $\log_{10}$ male volume | **Mating system** | 0.96 | 0.44 | 0.05 | $F_{2,98} = 3.47$ | **0.035** | Search – Guard = -0.12 ± 0.08, p = 0.24<br>Search – Mono = 0.22 ± 0.14, p = 0.24<br>**Guard – Mono = 0.34 ± 0.14, p = 0.04** |
| | $\log_{10}$ SSD | **Mating system** | 0.60 | 0.38 | 0.29 | $F_{2,98} = 21.36$ | <.0001 | **Search – Guard = 0.30 ± 0.05, p <.0001**<br>Search – Mono = -0.05 ± 0.09, p = 0.54<br>**Guard – Mono = -0.36 ± 0.09, p = 0.0002** |
| 1 | $\log_{10}$ female volume | **Flight dimorphism** | 0.98 | 0.58 | 0.09 | $F_{2,193} = 10.6$ | <.0001 | **Male only – None = 0.31 ± 0.07, p <.0001**<br>Male only – Both = 0.22 ± 0.11, p = 0.08<br>None – Both = -0.09 ± 0.09, p = 0.34 |
| | $\log_{10}$ male volume | **Flight dimorphism** | 0.98 | 0.52 | 0.04 | $F_{2,193} = 5.17$ | **0.006** | **Male only – None = 0.21 ± 0.07, p = 0.005**<br>Male only – Both = 0.18 ± 0.10, p = 0.15<br>None – Both = -0.02 ± 0.091, p = 0.77 |
| | $\log_{10}$ SSD | **Flight dimorphism** | 0.52 | 0.21 | 0.05 | $F_{2,193} = 5.95$ | **0.003** | **Male only – None = 0.17 ± 0.05, p = 0.002**<br>Male only – Both = 0.10 ± 0.08, p = 0.50<br>None – Both = -0.08 ± 0.08, p = 0.50 |
| 1 | $\log_{10}$ female volume | **Habitat** | 0.97 | 0.62 | 0.16 | $F_{2,192} = 19.0$ | <.0001 | Ground – Shrub = -0.11 ± 0.06, p = 0.06<br>**Ground – Canopy = -0.55 ± 0.09, p <.0001**<br>**Shrub – Canopy = -0.44 ± 0.08, p <.0001** |
| | $\log_{10}$ male volume | **Habitat** | 0.98 | 0.54 | 0.08 | $F_{2,192} = 9.67$ | **0.0001** | Ground – Shrub = -0.02 ± 0.06, p = 0.69<br>**Ground – Canopy = -0.37 ± 0.09, p = 0.0001**<br>**Shrub – Canopy = -0.35 ± 0.08, p = 0.0001** |
| | $\log_{10}$ SSD | **Habitat** | 0.52 | 0.23 | 0.07 | $F_{2,192} = 8.62$ | **0.0003** | **Ground – Shrub = -0.09 ± 0.05, p = 0.05**<br>**Ground – Canopy = -0.26 ± 0.06, p = 0.0002**<br>**Shrub – Canopy = -0.16 ± 0.06, p = 0.01** |
| 5 | $\log_{10}$ female volume | **$\log_{10}$ Female reproductive output** | 0.31 | 0.82 | 0.76 | $F_{1,62} = 137.3$ | <.0001 | **0.44 ± 0.04** |
| | | Climate PC1 | | | | $F_{1,62} = 1.76$ | 0.19 | -0.05 ± 0.04 |
| | | Mating system | | | | $F_{2,62} = 2.69$ | 0.08 | Search – Guard = 0.03 ± 0.06, p = 0.61<br>Search – Mono = 0.22 ± 0.10, p = 0.08<br>Guard – Mono = 0.19 ± 0.09, p = 0.08 |
| | | Flight dimorphism | | | | $F_{2,62} = 0.95$ | 0.39 | Male only – None = -0.01 ± 0.08, p = 0.89<br>Male only – Both = -0.14 ± 0.11, p = 0.58<br>None – Both = -0.13 ± 0.10, p = 0.58 |
| | | Habitat | | | | $F_{2,62} = 1.07$ | 0.35 | Ground – Shrub = -0.05 ± 0.07, p = 0.60<br>Ground – Canopy = -0.14 ± 0.09, p = 0.45<br>Shrub – Canopy = -0.09 ± 0.08, p = 0.59 |
| 5 | $\log_{10}$ male volume | **$\log_{10}$ Female reproductive output** | 0.16 | 0.75 | 0.71 | $F_{1,62} = 104.5$ | <.0001 | **0.43 ± 0.04** |
| | | Climate PC1 | | | | $F_{1,62} = 0.08$ | 0.78 | -0.01 ± 0.04 |
| | | **Mating system** | | | | $F_{2,62} = 10.9$ | **0.0001** | Search – Guard = -0.13 ± 0.07, p = 0.09<br>**Search – Mono = 0.34 ± 0.11, p = 0.006**<br>**Guard – Mono = 0.47 ± 0.10, p = 0.0001** |
| | | Flight dimorphism | | | | $F_{2,62} = 2.90$ | 0.06 | Male only – None = -0.17 ± 0.09, p = 0.14<br>Male only – Both = -0.24 ± 0.12, p = 0.14<br>None – Both = -0.07 ± 0.12, p = 0.56 |
| | | Habitat | | | | $F_{2,62} = 0.12$ | 0.88 | Ground – Shrub = 0.009 ± 0.09, p = 1<br>Ground – Canopy = -0.037 ± 0.11, p = 1<br>Shrub – Canopy = -0.046 ± 0.09, p = 1 |
| 5 | $\log_{10}$ SSD | $\log_{10}$ Female reproductive output | 0.26 | 0.43 | 0.40 | $F_{1,62} = 0.06$ | 0.81 | 0.009 ± 0.04 |
| | | Climate PC1 | | | | $F_{1,62} = 2.03$ | 0.16 | -0.055 ± 0.04 |
| | | **Mating system** | | | | $F_{2,62} = 9.83$ | **0.0002** | **Search – Guard = 0.19 ± 0.06, p = 0.009**<br>Search – Mono = -0.16 ± 0.10, p = 0.10<br>**Guard – Mono = -0.35 ± 0.09, p = 0.0007** |
| | | **Flight dimorphism** | | | | $F_{2,62} = 3.38$ | **0.04** | **Male only – None = 0.20 ± 0.08, p = 0.04**<br>Male only – Both = 0.15 ± 0.10, p = 0.31<br>None – Both = -0.04 ± 0.10, p = 0.66 |
| | | Habitat | | | | $F_{2,62} = 1.99$ | 0.15 | Ground – Shrub = -0.13 ± 0.07, p = 0.23<br>Ground – Canopy = -0.17 ± 0.09, p = 0.23<br>Shrub – Canopy = -0.04 ± 0.08, p = 0.62 |

**Table S3:** Ultimate predictors of variation in female and male body elongation (PC1) and SShD_elongation_. The table presents results of single and multiple predictor PGLS models. The most likely value of Pagel’s lambda (phylogenetic signal) is presented along with ANOVA outputs (using marginal sum of squares obtained by deleting a term from the model at a time) and either estimated effect sizes or post-hoc pairwise comparisons between estimated marginal means using the Holm method to account for multiple testing, respectively for continuous or categorical explanatory variables. The proportions of the variance explained by the full model R^2^_full_ (phylogeny + fixed effects) and explained by fixed effects alone R^2^_fixed_ (after accounting for phylogeny) are also reported.

| # predictors | Response | Predictor | $\lambda$ | $R^2_{full}$ | $R^2_{fixed}$ | $F_{df1,df2}$ | P | Effect size ± SE (continuous) OR Post-hoc pairwise tests (Holm) (categorical) |
| --- | --- | --- | --- | --- | --- | --- | --- | --- |
| 1 | Female PC1 | log<sub>10</sub> Female reproductive output | 0.95 | 0.47 | 0.01 | $F_{1,85} = 2.14$ | 0.14 | -0.36 ± 0.25 |
| | Male PC1 | log<sub>10</sub> Female reproductive output | 0.96 | 0.50 | 0.03 | $F_{1,85} = 3.34$ | 0.07 | -0.49 ± 0.27 |
| | SShD<sub>elongation</sub> | log<sub>10</sub> Female reproductive output | 0.93 | 0.33 | 0 | $F_{1,85} = 0.20$ | 0.65 | -0.06 ± 0.12 |
| 1 | Female PC1 | Climate PC1 | 0.96 | 0.59 | 0 | $F_{1,184} = 1.10$ | 0.29 | -0.09 ± 0.09 |
| | Male PC1 | Climate PC1 | 0.96 | 0.63 | 0 | $F_{1,184} = 0.34$ | 0.56 | -0.05 ± 0.09 |
| | SShD<sub>elongation</sub> | Climate PC1 | 0.75 | 0.29 | 0 | $F_{1,184} = 0.96$ | 0.33 | 0.04 ± 0.04 |
| 1 | Female PC1 | **Mating system** | 1 | 0.57 | 0.36 | $F_{2,95} = 27.7$ | <.0001 | Search – Guard = 2.74 ± 0.38, p <.0001<br>Search – Mono = 3.22 ± 0.73, p = 0.0001<br>Guard – Mono = 0.48 ± 0.69, p = 0.48 |
| | Male PC1 | **Mating system** | 0.97 | 0.65 | 0.32 | $F_{2,95} = 23.4$ | <.0001 | Search – Guard = 2.65 ± 0.43, p <.0001<br>Search – Mono = 4.02 ± 0.78, p <.0001<br>Guard – Mono = 1.37 ± 0.73, p = 0.06 |
| | SShD<sub>elongation</sub> | **Mating system** | 0.88 | 0.35 | 0.06 | $F_{2,95} = 3.96$ | 0.02 | Search – Guard = 0.25 ± 0.22, p = 0.28<br>Search – Mono = 1.11 ± 0.40, p = 0.02<br>Guard – Mono = 0.87 ± 0.38, p = 0.05 |
| 1 | Female PC1 | **Flight dimorphism** | 0.99 | 0.59 | 0.07 | $F_{2,183} = 7.82$ | 0.0006 | Male only – None = 0.40 ± 0.41, p = 0.32<br>Male only – Both = -1.77 ± 0.63, p = 0.01<br>None – Both = -2.17 ± 0.55, p = 0.0003 |
| | Male PC1 | **Flight dimorphism** | 0.97 | 0.64 | 0.03 | $F_{2,183} = 3.56$ | 0.03 | Male only – None = 0.70 ± 0.43, p = 0.20<br>Male only – Both = -0.78 ± 0.69, p = 0.26<br>None – Both = -1.49 ± 0.63, p = 0.06 |
| | SShD<sub>elongation</sub> | Flight dimorphism | 0.75 | 0.30 | 0.02 | $F_{2,183} = 2.67$ | 0.07 | Male only – None = 0.25 ± 0.19, p = 0.24<br>Male only – Both = 0.72 ± 0.32, p = 0.07<br>None – Both = 0.47 ± 0.30, p = 0.24 |
| 1 | Female PC1 | **Habitat** | 0.97 | 0.65 | 0.14 | $F_{2,182} = 16.48$ | <.0001 | Ground – Shrub = -1.95 ± 0.34, p <.0001<br>Ground – Canopy = -1.62 ± 0.54, p = 0.006<br>Shrub – Canopy = 0.33 ± 0.50, p = 0.52 |
| | Male PC1 | **Habitat** | 0.96 | 0.68 | 0.12 | $F_{2,182} = 13.40$ | <.0001 | Ground – Shrub = -1.84 ± 0.36, p <.0001<br>Ground – Canopy = -1.77 ± 0.56, p = 0.004<br>Shrub – Canopy = 0.07 ± 0.53, p = 0.89 |
| | SShD<sub>elongation</sub> | Habitat | 0.76 | 0.29 | 0 | $F_{2,182} = 0.82$ | 0.44 | Ground – Shrub = 0.03 ± 0.18, p = 0.88<br>Ground – Canopy = -0.27 ± 0.25, p = 0.63<br>Shrub – Canopy = -0.30 ± 0.24, p = 0.63 |
| 5 | Female PC1 | log<sub>10</sub> Female reproductive output | 1 | 0.55 | 0.45 | $F_{1,60} = 0.51$ | 0.48 | -0.20 ± 0.28 |
| | | Climate PC1 | | | | $F_{1,60} = 1.88$ | 0.18 | -0.21 ± 0.15 |
| | | **Mating system** | | | | $F_{2,60} = 3.27$ | 0.04 | Search – Guard = 1.28 ± 0.57, p = 0.08<br>Search – Mono = 1.93 ± 0.85, p = 0.08<br>Guard – Mono = 0.66 ± 0.70, p = 0.36 |
| | | Flight dimorphism | | | | $F_{2,60} = 1.07$ | 0.35 | Male only – None = 0.15 ± 0.71, p = 0.84<br>Male only – Both = -1.14 ± 0.95, p = 0.48<br>None – Both = -1.28 ± 0.89, p = 0.47 |
| | | Habitat | | | | $F_{2,60} = 1.75$ | 0.18 | Ground – Shrub = -0.62 ± 0.57, p = 0.40<br>Ground – Canopy = -1.56 ± 0.83, p = 0.20<br>Shrub – Canopy = -0.94 ± 0.72, p = 0.40 |
| 5 | Male PC1 | log<sub>10</sub> Female reproductive output | 0.97 | 0.59 | 0.34 | $F_{1,60} = 0.51$ | 0.48 | -0.24 ± 0.34 |
| | | Climate PC1 | | | | $F_{1,60} = 3.40$ | 0.07 | -0.33 ± 0.18 |
| | | **Mating system** | | | | $F_{2,60} = 7.78$ | 0.001 | Search – Guard = 2.04 ± 0.64, p = 0.005<br>Search – Mono = 3.49 ± 0.95, p = 0.015<br>Guard – Mono = 1.45 ± 0.81, p = 0.08 |
| | | Flight dimorphism | | | | $F_{2,60} = 0.07$ | 0.93 | Male only – None = 0.09 ± 0.79, p = 1<br>Male only – Both = -0.30 ± 1.07, p = 1<br>None – Both = -0.39 ± 1.02, p = 1 |
| | | Habitat | | | | $F_{2,60} = 1.35$ | 0.27 | Ground – Shrub = -0.57 ± 0.65, p = 0.47<br>Ground – Canopy = -1.54 ± 0.94, p = 0.32<br>Shrub – Canopy = -0.97 ± 0.81, p = 0.47 |
| 5 | SShD<sub>elongation</sub> | log<sub>10</sub> Female reproductive output | 0.93 | 0.37 | 0.09 | $F_{1,60} = 0.32$ | 0.57 | -0.10 ± 0.17 |
| | | Climate PC1 | | | | $F_{1,60} = 0.34$ | 0.56 | -0.05 ± 0.08 |
| | | **Mating system** | | | | $F_{2,60} = 5.55$ | 0.006 | Search – Guard = 0.72 ± 0.32, p = 0.05<br>Search – Mono = 1.53 ± 0.47, p = 0.006<br>Guard – Mono = 0.81 ± 0.41, p = 0.05 |
| | | Flight dimorphism | | | | $F_{2,60} = 1.61$ | 0.21 | Male only – None = -0.05 ± 0.39, p = 0.90<br>Male only – Both = 0.83 ± 0.53, p = 0.27<br>None – Both = 0.88 ± 0.51, p = 0.27 |
| | | Habitat | | | | $F_{2,60} = 0.07$ | 0.94 | Ground – Shrub = 0.08 ± 0.33, p = 1<br>Ground – Canopy = -0.04 ± 0.47, p = 1<br>Shrub – Canopy = -0.13 ± 0.40, p = 1 |

**Table S4:** Ultimate predictors of variation in female and male relative wing size (PC2) and SShD_wings_. The table presents results of single and multiple predictor PGLS models. The most likely value of Pagel’s lambda (phylogenetic signal) is presented along with ANOVA outputs (using marginal sum of squares obtained by deleting a term from the model at a time) and either estimated effect sizes or post-hoc pairwise comparisons between estimated marginal means using the Holm method to account for multiple testing, respectively for continuous or categorical explanatory variables. The proportions of the variance explained by the full model R^2^_full_ (phylogeny + fixed effects) and explained by fixed effects alone R^2^_fixed_ (after accounting for phylogeny) are also reported.

| # predictors | Response | Predictor | $\lambda$ | $R^2_{full}$ | $R^2_{fixed}$ | $F_{df1,df2}$ | P | Effect size ± SE (continuous)<br>Post-hoc pairwise tests (Holm) (categorical) |
| --- | --- | --- | --- | --- | --- | --- | --- | --- |
| 1 | Female PC2 | $\log_{10}$ Female reproductive output | 0.69 | 0.24 | 0.02 | $F_{1,85} = 2.65$ | 0.11 | -0.26 ± 0.16 |
| | Male PC2 | $\log_{10}$ Female reproductive output | 0.78 | 0.34 | 0 | $F_{1,85} = 1.28$ | 0.26 | 0.24 ± 0.21 |
|  | SShD<sub>wings</sub> | **$\log_{10}$ Female reproductive output** | 0.84 | 0.42 | 0.11 | **$F_{1,85} = 12.1$** | **0.0008** | **0.51 ± 0.15** |
| 1 | Female PC2 | Climate PC1 | 0.83 | 0.37 | 0 | $F_{1,184} = 0.66$ | 0.42 | -0.04 ± 0.05 |
| | Male PC2 | Climate PC1 | 0.88 | 0.47 | 0 | $F_{1,184} = 0.85$ | 0.36 | 0.05 ± 0.05 |
| | SShD<sub>wings</sub> | Climate PC1 | 1 | 0.31 | 0.01 | $F_{1,184} = 3.39$ | 0.07 | 0.08 ± 0.04 |
| 1 | Female PC2 | Mating system | 0.69 | 0.22 | 0 | $F_{2,95} = 1.04$ | 0.36 | Search – Guard = 0.43 ± 0.30, p = 0.46<br>Search – Mono = 0.21 ± 0.52, p = 1<br>Guard – Mono = -0.22 ± 0.51, p = 1 |
|  | Male PC2 | **Mating system** | 0.82 | 0.31 | 0.08 | **$F_{2,95} = 5.25$** | **0.007** | **Search – Guard = 1.01 ± 0.35, p = 0.01**<br>**Search – Mono = 1.47 ± 0.62, p = 0.04**<br>**Guard – Mono = 0.46 ± 0.59, p = 0.44** |
|  | SShD<sub>wings</sub> | **Mating system** | 1 | 0.29 | 0.06 | **$F_{2,95} = 4.29$** | **0.02** | **Search – Guard = 0.49 ± 0.21, p = 0.04**<br>**Search – Mono = 1.01 ± 0.40, p = 0.04**<br>**Guard – Mono = 0.51 ± 0.38, p = 0.18** |
| 1 | Female PC2 | **Flight dimorphism** | 0.83 | 0.84 | 0.75 | **$F_{2,183} = 282.1$** | **<.0001** | **Male only – None = 0.59 ± 0.11, p <.0001**<br>**Male only – Both = -3.49 ± 0.18, p <.0001**<br>**None – Both = -4.08 ± 0.17, p <.0001** |
|  | Male PC2 | **Flight dimorphism** | 0.95 | 0.88 | 0.78 | **$F_{2,183} = 330.4$** | **<.0001** | **Male only – None = 2.50 ± 0.12, p <.0001**<br>**Male only – Both = -1.21 ± 0.20, p <.0001**<br>**None – Both = -3.71 ± 0.19, p <.0001** |
|  | SShD<sub>wings</sub> | **Flight dimorphism** | 0.84 | 0.70 | 0.54 | **$F_{2,183} = 110.2$** | **<.0001** | **Male only – None = 1.97 ± 0.14, p <.0001**<br>**Male only – Both = 2.41 ± 0.24, p <.0001**<br>**None – Both = 0.44 ± 0.22, p = 0.05** |
| 1 | Female PC2 | **Habitat** | 0.84 | 0.45 | 0.12 | **$F_{2,182} = 13.92$** | **<.0001** | **Ground – Shrub = -0.75 ± 0.19, p = 0.0002**<br>**Ground – Canopy = -1.33 ± 0.27, p <.0001**<br>**Shrub – Canopy = -0.58 ± 0.26, p = 0.03** |
|  | Male PC2 | **Habitat** | 0.86 | 0.62 | 0.27 | **$F_{2,182} = 35.7$** | **<.0001** | **Ground – Shrub = -1.01 ± 0.21, p <.0001**<br>**Ground – Canopy = -2.49 ± 0.30, p <.0001**<br>**Shrub – Canopy = -1.49 ± 0.28, p <.0001** |
|  | SShD<sub>wings</sub> | **Habitat** | 1 | 0.37 | 0.09 | **$F_{2,182} = 9.96$** | **0.0001** | **Ground – Shrub = -0.17 ± 0.16, p = 0.29**<br>**Ground – Canopy = -1.21 ± 0.27, p = 0.0001**<br>**Shrub – Canopy = -1.04 ± 0.26, p = 0.0002** |
| 5 | Female PC2 | $\log_{10}$ Female reproductive output | 1 | 0.86 | 0.93 | $F_{1,60} = 0.08$ | 0.77 | 0.02 ± 0.08 |
|  |  | Climate PC1 |  |  |  | **$F_{1,60} = 4.89$** | **0.03** | **-0.10 ± 0.04** |
| | | Mating system | | | | $F_{2,60} = 0.09$ | 0.91 | Search – Guard = 0.06 ± 0.17, p = 1<br>Search – Mono = 0.01 ± 0.25, p = 1<br>Guard – Mono = -0.05 ± 0.21, p = 1 |
|  |  | **Flight dimorphism** |  |  |  | **$F_{2,60} = 132.1$** | **<.0001** | **Male only – None = 0.68 ± 0.21, p = 0.002**<br>**Male only – Both = -3.54 ± 0.28, p <.0001**<br>**None – Both = -4.22 ± 0.26, p <.0001** |
| | | Habitat | | | | $F_{2,60} = 0.15$ | 0.86 | Ground – Shrub = -0.02 ± 0.17, p = 1<br>Ground – Canopy = 0.09 ± 0.24, p = 1<br>Shrub – Canopy = 0.11 ± 0.21, p = 1 |
| 5 | Male PC2 | $\log_{10}$ Female reproductive output | 0.99 | 0.82 | 0.87 | $F_{1,60} = 0.59$ | 0.44 | -0.09 ± 0.12 |
| | | Climate PC1 | | | | $F_{1,60} = 1.05$ | 0.31 | -0.07 ± 0.07 |
| | | Mating system | | | | $F_{2,60} = 0.14$ | 0.87 | Search – Guard = 0.08 ± 0.24, p = 1<br>Search – Mono = 0.19 ± 0.36, p = 1<br>Guard – Mono = 0.11 ± 0.30, p = 1 |
|  |  | **Flight dimorphism** |  |  |  | **$F_{2,60} = 72.2$** | **<.0001** | **Male only – None = 2.75 ± 0.31, p <.0001**<br>**Male only – Both = -1.25 ± 0.41, p = 0.003**<br>**None – Both = -4.0 ± 0.38, p <.0001** |
| | | Habitat | | | | $F_{2,60} = 0.64$ | 0.53 | Ground – Shrub = 0.001 ± 0.24, p = 1<br>Ground – Canopy = -0.34 ± 0.36, p = 0.81<br>Shrub – Canopy = -0.34 ± 0.31, p = 0.81 |
| 5 | SShD<sub>wings</sub> | $\log_{10}$ Female reproductive output | 0.38 | 0.68 | 0.63 | $F_{1,60} = 0.17$ | 0.68 | 0.06 ± 0.14 |
| | | Climate PC1 | | | | $F_{1,60} = 0.32$ | 0.58 | 0.03 ± 0.06 |
| | | Mating system | | | | $F_{2,60} = 0.05$ | 0.95 | Search – Guard = -0.03 ± 0.24, p = 1<br>Search – Mono = 0.08 ± 0.36, p = 1<br>Guard – Mono = 0.11 ± 0.34, p = 1 |
|  |  | **Flight dimorphism** |  |  |  | **$F_{2,60} = 32.5$** | **<.0001** | **Male only – None = 2.05 ± 0.29, p <.0001**<br>**Male only – Both = 2.54 ± 0.39, p <.0001**<br>**None – Both = 0.49 ± 0.39, p = 0.21** |
| | | Habitat | | | | $F_{2,60} = 0.47$ | 0.63 | Ground – Shrub = -0.08 ± 0.28, p = 1<br>Ground – Canopy = -0.34 ± 0.37, p = 1<br>Shrub – Canopy = -0.26 ± 0.31, p = 1 |

